# The intestinal microbiome shapes responses to the GD2/GD3 cancer vaccine in children with high-risk neuroblastoma

**DOI:** 10.64898/2026.11.16.766479

**Authors:** Lanying Ma, Kimon Argyropoulos, Kanaka Dhuri, Nicholas R. Waters, Teng Fei, Irene Y. Cheung, Xiaohong Wu, Yunlin Zhang, Leena Babiker, Katlyn Lederer, Leana Harford, Nadia Brijmohan, Chi Nguyen, Kristen Victor, Jianmin Yang, Anqi Dai, Jonathan U. Peled, Alan Hanash, Stephen S. Roberts, Jaap Jan Boelens, Stefan Nierkens, Gretchen Diehl, Kate A. Markey, Ying Taur, Marina Burgos da Silva, Brian H. Kushner, Ivan Maillard, Govind Ragupathi, Nai-Kong Cheung, Marcel van den Brink, Shakeel Modak, Oriana Miltiadous

## Abstract

The intestinal microbiome influences vaccine responses, but its role in therapeutic cancer vaccination remains largely unexplored. We investigated the gut microbiome, vaccine immunogenicity and clinical outcomes in 137 children with high-risk neuroblastoma receiving GD2/GD3 vaccination with oral β-glucan. Microbiome-perturbing antibiotic exposure was associated with inferior progression-free survival (PFS). Among patients vaccinated in first remission, higher α-diversity and enrichment of *Lachnospiraceae*, including *Blautia*, were associated with higher anti-GD2 IgG1 responses, while higher *Clostridia* abundance was associated with superior PFS. Patients had microbiome profiles resembling those of chronologically younger healthy children, consistent with delayed microbiome maturation. A greater discrepancy between patients’ chronological age and the age of microbiome-matched healthy children was associated with inferior PFS among patients in first remission. In a complementary mouse model, antibiotic treatment impaired vaccine-induced anti-GD2 IgG responses and accelerated tumor growth. Together, these findings support a role for the intestinal microbiome in therapeutic cancer vaccine responses and identify microbiome maturation as a clinically relevant feature in pediatrics.

## Introduction

High-risk neuroblastoma (HR-NB) is an aggressive pediatric malignancy that remains challenging to treat. Despite intensive multimodal therapy, it is associated with substantial morbidity and mortality due to both disease progression and treatment-related toxicities.^1,2^ Neuroblastoma tumors express high levels of the cell-surface ganglioside 2 (GD2), a glycosphingolipid that serves as an established target for monoclonal antibody-based immunotherapy.^3–5^ Anti-GD2 monoclonal antibody therapy has significantly improved outcomes in children with HR-NB.^6–9^ However, even with the addition of anti-GD2 monoclonal antibody treatment, approximately 55% of patients with HR-NB still relapse.^6,9,10^ To improve the durability of remission, a vaccine targeting the neuroblastoma-associated gangliosides GD2 and GD3, conjugated to the immunogenic carrier protein keyhole limpet hemocyanin (KLH) along with the saponin adjuvant QS-21, has been evaluated and continues to be investigated in clinical trials in patients who achieve remission.^11,12^ β-glucan, an immunomodulatory polysaccharide, is administered orally alongside this vaccine and has been shown to enhance antitumor immune responses and induce trained immunity.^13–18^

In a clinical trial of 102 patients with HR-NB, administration of GD2/GD3 vaccine in combination with oral β-glucan elicited immunoglobulin G1 (IgG1) responses, with higher anti-GD2 IgG1 titers associated with superior progression-free survival (PFS).^11^ In a subsequent randomized controlled trial of 107 patients, earlier β-glucan initiation was associated with higher anti-GD2 IgG1 titers at weeks 20, 32 and 52, whereas titers at week 8 were similar between groups. Higher week 8 anti-GD2 IgG1 titers remained significantly associated with superior PFS.^19^ Importantly, the two groups did not differ in PFS or in the proportion of patients who remained seronegative throughout the vaccine course, defined as anti-GD2 IgG1 titers below the assay limit of detection at all measured time points.

Several lines of evidence implicate the intestinal microbiome as a key modulator of vaccine responses.^20–25^ Germ-free mice and TLR5-deficient mice with impaired sensing of bacterial flagellin exhibit diminished immune responses to some vaccines.^22,23^ Similarly, early-life antibiotic exposure reduces antibody titer responses to childhood vaccines in mice, a phenotype that can be rescued by fecal microbiota transplant (FMT).^24^ In a randomized clinical trial in healthy adults, antibiotic exposure impaired antibody responses to influenza H1N1 vaccine among individuals with low pre-existing immunity, but not among those with high pre-existing immunity.^25^ The intestinal microbiota is also an important modulator of responses to cancer immunotherapy. In adult patients receiving immune checkpoint inhibitors or CAR T-cell therapy, antibiotic exposures are associated with worse responses, while enrichment of commensal taxa including *Lachnospiraceae* and *Clostridiales* is associated with improved PFS.^26–30^ Preclinical studies support a causal role for the gut microbiome in shaping cancer immunotherapy responses,^31–33^ while FMT has shown the potential to enhance responses in both preclinical models and early-phase clinical trials.^34–36^ However, whether the intestinal microbiome influences responses to therapeutic cancer vaccines in patients remains largely unexplored.

Unlike adults, children have a dynamically developing intestinal microbiome that undergoes maturation alongside the immune system during early childhood.^37–47^ Children with HR-NB are exposed to repeated courses of chemotherapy, radiation, and broad-spectrum antibiotics, with a subset also undergoing stem cell transplantation, all of which can profoundly alter the intestinal microbiota.^48–51^ Despite these developmental and treatment-related factors, the role of the intestinal microbiome in shaping responses to cancer immunotherapy in children remains poorly understood.

Here, we hypothesized that the intestinal microbiome influences responses to GD2/GD3 vaccination in children with HR-NB. We investigated associations between the microbiome, vaccine immunogenicity and clinical outcomes, compared microbiome composition with that of healthy children to evaluate age-related microbiome features, and used a preclinical model of antibiotic-mediated microbiome perturbation to experimentally investigate its effects on vaccine responses.

## Results

### Patient characteristics

To investigate the relationship between the intestinal microbiome and responses to GD2/GD3 vaccination, we collected stool samples longitudinally throughout the vaccine course in a cohort of 137 patients (NCT00911560; **Fig. 1a**). Because participants were enrolled in different phases of the clinical trial (**Extended Data Fig. 1a**), we first examined the protocol-defined clinical trial outcomes relevant to the cohorts represented in the microbiome study (**Suppl. Table 1**).

**Figure 1.**
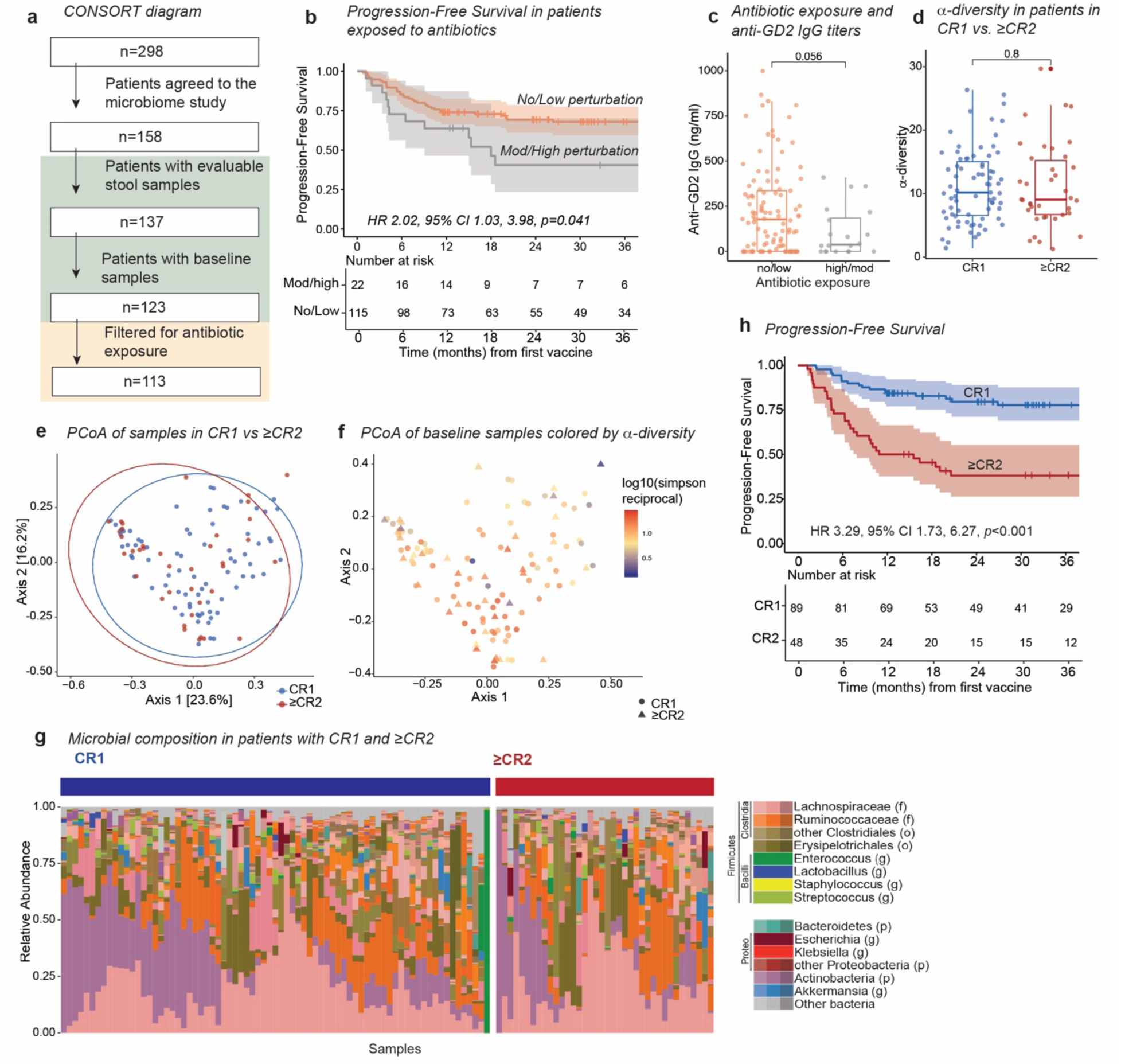
Antibiotic exposure is associated with inferior progression-free survival in patients receiving GD2/GD3 vaccination. (**a**) CONSORT diagram showing patient enrollment and sample availability. (**b**) Kaplan-Meier analysis of progression-free survival in patients exposed to moderate/high microbiome-perturbing antibiotics compared with patients receiving no antibiotics or low microbiome-perturbing antibiotics. (**c**) Anti-GD2 IgG titers in patients exposed to moderate/high microbiome-perturbing antibiotics compared with patients receiving no antibiotics or low microbiome-perturbing antibiotics. (**d-g**) Comparison of baseline gut microbiome features between patients vaccinated in CR1 or ≥CR2. (**d**) α-diversity at baseline. (**e,f**) PCoA of baseline microbial community composition. (**g**) Baseline microbial composition in patients in CR1 and ≥CR2. (**h**) Kaplan-Meier analysis of PFS according to remission status (CR1 versus ≥CR2). Hazard ratios and *p* values are derived from a multivariable Cox regression model. Statistical comparisons in **c,d** were performed using two-sided Wilcoxon rank-sum tests.

Baseline clinical characteristics by clinical trial subcohort are shown in **Suppl. Table 2**. Among the 24 microbiome-study participants enrolled in the Phase II cohort, 14 were in CR1 and 10 were in ≥CR2. One-year relapse-free survival in the CR1 cohort was 92.9% (95% CI, 82.0-100%; **Extended Data Fig. 1b**). Of the 10 patients in ≥CR2, 2 experienced relapse by 6 months and 8 remained relapse-free at 6 months and (80%; **Extended Data Fig. 1c**). Among the 104 microbiome-study participants enrolled during the Phase II randomization cohort, 73 were randomized to either initiate β-glucan at week 6 (n=35; Group 1) or at week 1 (n=38; Group 2), and 31 were enrolled in the nonrandomized Group 3 exploratory cohort (β-glucan initiated at week 6). Consistent with the previously reported trial results, week 32 anti-GD2 antibody titers were higher among patients who initiated glucan at week 1 than among those who initiated glucan at week 6 after adjustment for CR status (*p*=0.013, linear regression; **Extended Data Fig. 1d**).^19^ We additionally evaluated week 8 anti-GD2 antibody titers, which were used in subsequent analyses in this study, and observed no significant difference between randomization groups at this time point, consistent with the previously reported analysis (*p*=0.30; **Extended Data Fig. 1e**).^19^

Study design and sample collection are shown in **Extended Data Fig 2a,b**. Gut microbiome profiling was performed on serial stool samples with 16S rRNA sequencing, with metagenomic shotgun sequencing performed on a subset of these samples. Of the 137 patients, 89 (65%) received the vaccine in first complete remission (CR1), whereas 48 (35%) had experienced one or more prior relapses before vaccination (≥CR2). Patients in the ≥CR2 group were older at vaccine initiation and were more likely to have undergone autologous stem cell transplantation (HCT) than those in CR1 (**Table 1**). Most patients in both groups had metastatic disease at initial diagnosis (97% and 98%, respectively), and *MYCN* amplification, an established prognostic feature in neuroblastoma^52^, was present in 33% of the overall cohort. Anti-GD2 IgG1 titers were measured at week 8 of the study, following the fourth vaccination, as previously described.^11^

**Table 1.**
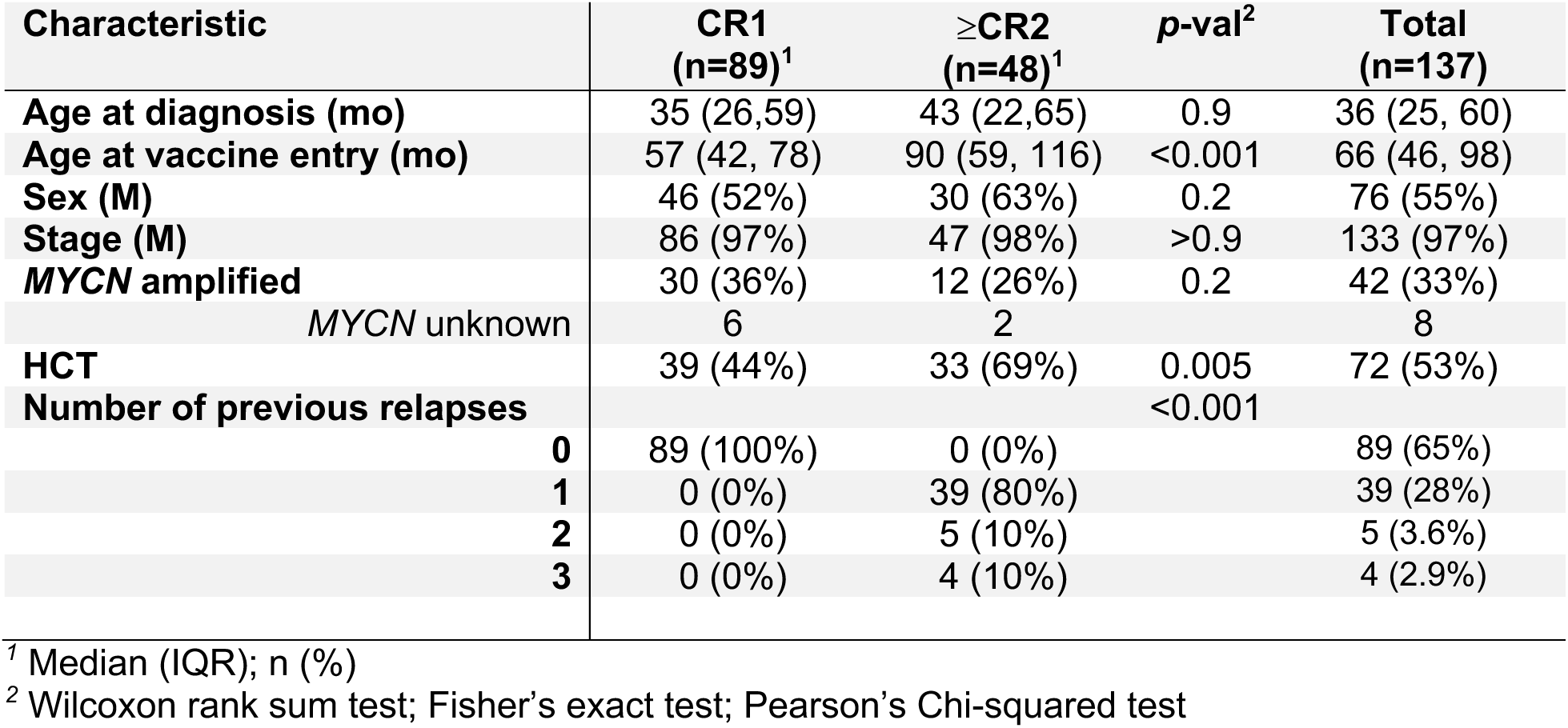
Patient characteristics.

### Antibiotic associations with outcomes in patients receiving the GD2/GD3 vaccine

To investigate the association between antibiotic exposure and the intestinal microbiota, we evaluated antibiotic use within 15 days before or after vaccine initiation. Samples were considered antibiotic-exposed if patients received at least one day of antibiotic therapy before sample collection. To assess the potential influence of antibiotics on fecal α-diversity, we compared paired samples collected before and after antibiotic treatment. Seven patients had paired samples available (ceftriaxone, n=5; metronidazole, n=1; amoxicillin/clavulanate, n=1), and α-diversity decreased following antibiotic exposure (Simpson reciprocal index, pre-exposure, 15.6; post-exposure, 10.4; *p*=0.04; **Extended Data Fig. 2c**).

Based on previously published effects on the intestinal microbiome, antibiotics were subsequently classified as having low or moderate/high microbiome-perturbing potential (**Suppl. Table 3**).^53–57^ Among patients exposed to moderate/high microbiome-perturbing antibiotics, most received antibiotics empirically for non-neutropenic fever (n=18; **Suppl. Table 4**). Only three patients had a documented infection, and two received prophylactic antibiotics for a procedure. None required intensive care admission, and most were managed in the outpatient setting (n=21/22, 95%). Exposure to moderate/high microbiome-perturbing antibiotics was associated with inferior PFS after adjustment for age, remission status (CR) and *MYCN* amplification (HR 2.02, 95% CI 1.03-3.98; *p*=0.041; **Fig. 1b, Suppl. Table 5**). Patients exposed to moderate/high microbiome-perturbing antibiotics had lower anti-GD2 IgG1 titers, although this difference did not reach statistical significance (median, 36.8 vs. 178.0 ng/mL; *p*=0.056; **Fig. 1c**).

The association between antibiotic exposure and clinical outcomes suggested a potential role for the gut microbiome in modulating responses to GD2/GD3 vaccination. We therefore hypothesized that greater gut microbiome diversity and enrichment of commensal anaerobes would be associated with higher anti-GD2 IgG1 titers and superior PFS. Given that antibiotic exposure can alter microbiome composition and was itself associated with clinical outcomes, we accounted for antibiotic exposure when selecting samples for downstream analyses. Samples collected before subsequent exposure to moderate/high microbiome-perturbing antibiotics did not capture the effects of that exposure on the microbiome and were therefore excluded from downstream microbiome analyses (**Extended Data Fig. 2d,e**). Application of this criterion resulted in 113 evaluable baseline samples from 113 patients for subsequent analyses.

### CR1 and ≥CR2 patients have similar baseline fecal microbiome composition

Given the differences in clinical characteristics between patients in CR1 and ≥CR2, we first examined whether baseline gut microbiome features differed between these groups (**Fig. 1d-g).** α-diversity was similar between CR1 and ≥CR2 patients (CR1: median=9.2; ≥CR2: median=10.2, *p*=0.8; **Fig. 1d**). Similarly, overall microbial composition did not differ significantly between the two groups (PERMANOVA, *p*=0.42, **Fig. 1e-g**). Because age and autologous HCT have previously been associated with alterations in the intestinal microbiome,^37,49^ we also examined microbiome composition according to these variables. Neither age (<5 vs. ≥5 years) nor transplant status was significantly associated with differences in microbiome composition (PERMANOVA, *p*=0.54 for age; *p*=0.068 for transplant status; **Extended Data Fig. 2f**).

Patients in ≥CR2 had significantly inferior PFS compared with patients in CR1 (HR 3.29, 95% CI 1.73-6.27; *p*<0.001, **Fig. 1h, Suppl. Table 5**). After a median follow-up of 32 months, 47 relapses and 10 deaths had occurred. Both relapses and deaths were more frequent in the ≥CR2 cohort, with 29 relapses and 7 deaths among 48 patients (60% and 15%, respectively), compared with 18 relapses and 3 deaths among 89 patients in CR1 (20% and 3%, respectively). All deaths were preceded by relapse. Given the marked differences in clinical outcomes between CR1 and ≥CR2 patients, subsequent analyses were stratified by remission status.

### Higher α-diversity and enrichment of Blautia and Lachnospiraceae are associated with higher anti-GD2 IgG1 titers in CR1 patients

We first examined the relationship between baseline fecal microbiota α-diversity and anti-GD2 IgG1 titers, measured at week 8 of the vaccine study. Among CR1 patients, baseline α-diversity was positively correlated with anti-GD2 IgG1 titers (Spearman, R=0.28, *p*=0.015), whereas no association was found in ≥CR2 patients (R=-0.16, *p*=0.41; **Fig. 2a**). We next investigated whether specific microbial communities were associated with anti-GD2 IgG1 titers by applying topic modeling to identify patterns of co-occurring taxa.^58^ This approach identified six distinct microbial community signatures among baseline samples from patients in CR1 and ≥CR2 (**Fig. 2b; Extended Data Fig. 3a-c**). Signatures were named according to the highest-weight taxon, with the two highest-weight taxa used to distinguish signatures sharing the same dominant taxon. Among patients in CR1, the six signatures were characterized by (1) *Blautia/Romboutsia*, (2) *Erysipelatoclostridium*, (3) *Faecalibacterium*, (4) *Bifidobacterium*, (5) *Blautia/Anaerostipes*, and (6) *Ruminococcus* (**Fig. 2b**). Among patients in ≥CR2, the six signatures were characterized by (1) *Ruminococcus*, (2) *Blautia/Faecalibacterium*, (3) *Romboutsia*, (4) *Blautia/Erysipelatoclostridium*, (5) *Bifidobacterium*, and (6) *Blautia/Anaerostipes* (**Extended Data Fig. 3a**).

**Figure 2.**
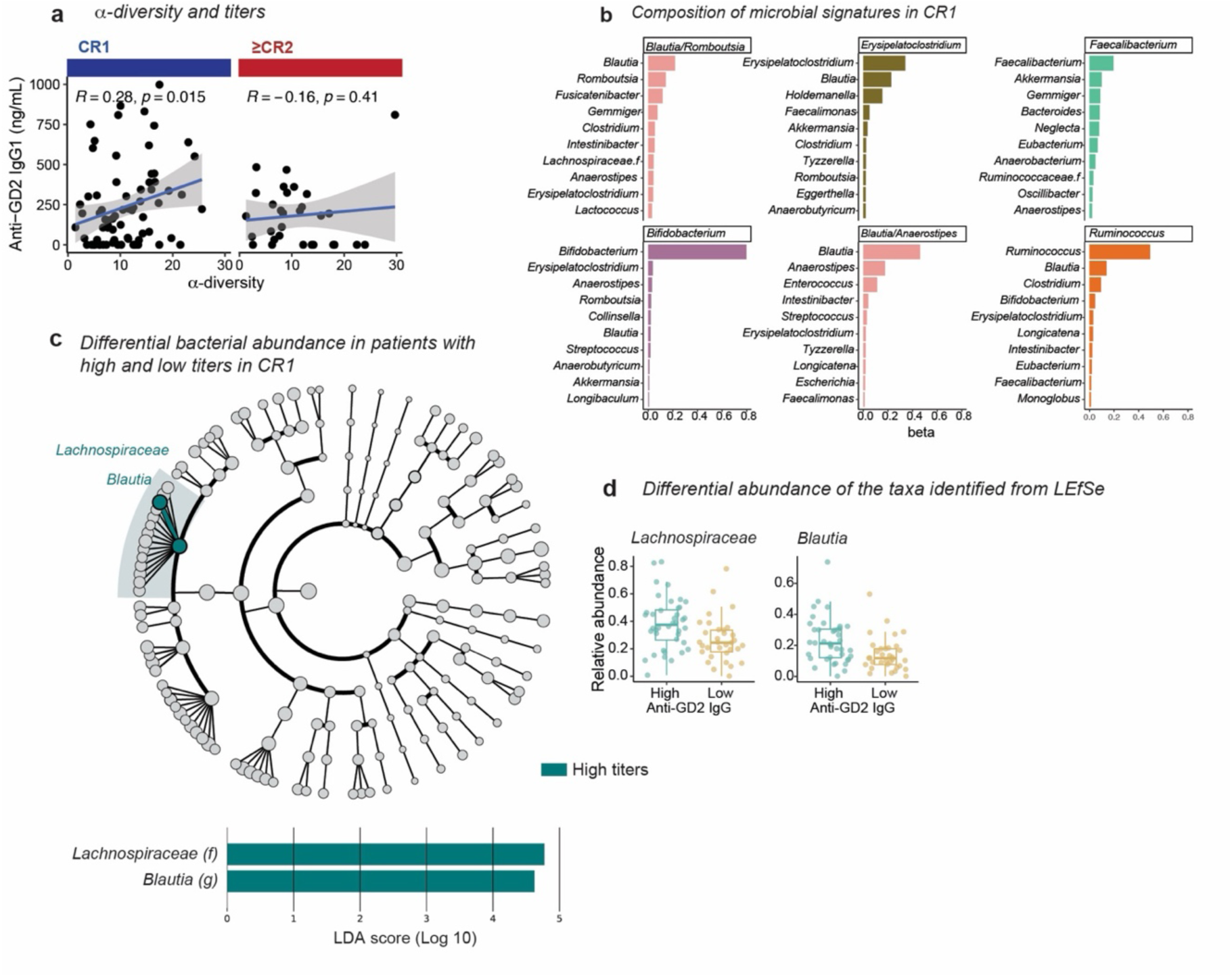
α-diversity and enrichment of *Lachnospiraceae* and *Blautia* are associated with higher anti-GD2 IgG1 in CR1 patients. (**a**) Association between baseline α-diversity and anti-GD2 IgG1 titers, stratified by CR status. (**b**) Microbial community signatures identified by topic modeling and labeled according to the dominant taxon within each signature for CR1. (**c**) LEfSe analysis identifying differentially abundant taxa between CR1 patients with high versus low anti-GD2 IgG1 titers. The analysis included 71 patients, of whom 38 had high and 33 had low titers. (**d**) Relative abundance of taxa identified by LEfSe as enriched in patients with high or low anti-GD2 IgG1 titers. Statistical comparisons in **a** were performed using Spearman correlation.

In a multivariable linear regression model of log-transformed anti-GD2 IgG1 titers, the *Blautia/Anaerostipes* signature showed a positive association with anti-GD2 IgG1 titers in CR1 patients after adjustment for *MYCN* amplification, HCT status, and α-diversity (**Suppl. Table 6,** β=1.21, *p=*0.004). No microbial community signature was significantly associated with anti-GD2 IgG1 titers among patients in ≥CR2 (**Suppl. Table 6**). To complement the topic modeling, we next used linear discriminant analysis effect size (LEfSe)^59^ to identify taxa associated with high versus low anti-GD2 IgG1 responses, using the previously reported cutoff of 150 ng/mL that was associated with differences in PFS.^11^ An FDR threshold of 0.05 was applied to account for multiple testing. Among CR1 patients, LEfSe identified enrichment of both the *Lachnospiraceae* family and the genus *Blautia* (a member of the *Lachnospiraceae* family) in patients with high anti-GD2 IgG1 titers (**Fig. 2c,d**). Taken together, these findings suggest that higher microbiome diversity and enrichment of taxa within the *Lachnospiraceae* family, including *Blautia*, are associated with superior humoral responses to GD2/GD3 vaccination.

### Clostridia abundance and an Erysipelatoclostridium signature are associated with PFS in CR1 patients

We next investigated whether baseline fecal microbiota features were associated with PFS. α-diversity was not associated with PFS in either CR1 (HR 0.95, 95% CI 0.86-1.04; *p*=0.28) or ≥CR2 patients (HR 1.02, 95% CI 0.96-1.10; *p*=0.52, **Suppl. Table 7**). We then evaluated microbial community signatures as predictors of PFS. Among patients in CR1, a higher abundance of the *Erysipelatoclostridium* signature was associated with inferior PFS in multivariable analysis adjusting for age, *MYCN* amplification, and the *Ruminococcus* signature (HR 20.73, 95% CI 2.05-209.58, *p*=0.01, **Fig. 3a, Suppl. Table 7**). Specifically, each 1-percentage-point increase in *Erysipelatoclostridium* signature abundance was associated with a 3.1% higher hazard of relapse. The wide confidence interval and high effect size may in part reflect the skewed distribution of the *Erysipelatoclostridium* signature (**Extended Data Fig. 3d**). No microbial community signature was significantly associated with PFS among patients in ≥CR2 (**Suppl. Table 7**).

**Figure 3.**
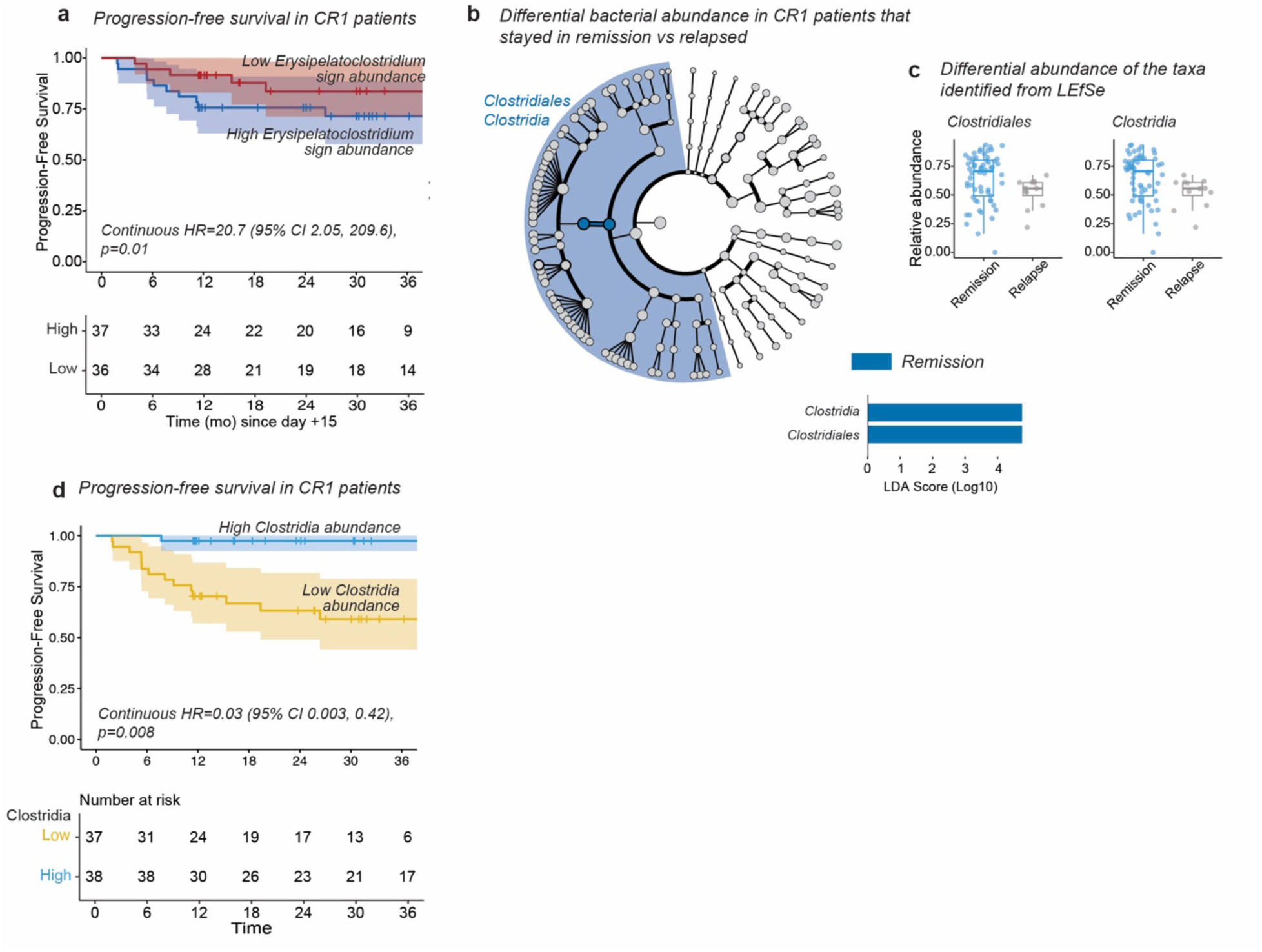
*Erysipelatoclostridium* signature abundance and *Clostridia* abundance are associated with progression-free survival in CR1 patients. (**a**) Kaplan-Meier analysis of progression-free survival according to the *Erysipelatoclostridium* topic signature among CR1 patients. (**b**) LEfSe analysis identifying differentially abundant taxa between CR1 patients who relapsed within 12 months of vaccine initiation and those who remained in remission. The analysis included 70 patients, of whom 11 relapsed and 59 remained in remission. (**c**) Relative abundance of taxa identified by LEfSe as differentially enriched according to 12-month relapse status. (**d**) Kaplan-Meier analysis of PFS in CR1 patients with high versus low *Clostridia* abundance, dichotomized at the median. Hazard ratio and *p* value are derived from separate multivariable Cox regression models in which *Clostridia* abundance and *Erysipelatoclostridium* topic signature were modeled as a continuous variable.

To identify individual taxa associated with PFS, we next used LEfSe to compare baseline microbiome composition between patients whose disease relapsed within 12 months and those who remained in remission; this time point captured 80% of observed relapses (**Extended Data Fig. 3e**). An FDR threshold of 0.05 was applied to account for multiple testing. Among CR1 patients, enrichment of the class *Clostridia* and order *Clostridiales* was observed in patients who remained in remission compared with those who relapsed (**Fig. 3b,c**). No significant associations were identified among ≥CR2 patients. Consistent with these findings, higher *Clostridia* abundance was independently associated with superior PFS in multivariable Cox regression, adjusting for age and *MYCN* amplification (HR 0.03, 95% CI 0.003-0.42, *p*=0.008, **Fig. 3d**, **Suppl. Table 8**). Specifically, each 1-percentage-point increase in *Clostridia* relative abundance was associated with a 3.3% lower hazard of relapse.

### Greater microbiome relative age gap is associated with inferior PFS in CR1

The intestinal microbiome undergoes substantial maturation during early childhood, characterized by increasing α-diversity and microbial complexity with age.^37–41^ Although much of this development occurs during infancy, the microbiome continues to mature throughout childhood and remains distinct from the adult microbiome even at 5 years of age, and undergoes further compositional changes during puberty.^38,41^ Because children with HR-NB are exposed to multiple microbiome-disrupting therapies, including chemotherapy, radiation, stem cell transplantation, and antibiotics,^48–51^ we investigated whether their fecal microbiome at vaccine initiation more closely resembled that of healthy children of similar chronological age or that of younger healthy children. Vaccine initiation occurred at least one month after completion of standard therapy. We hypothesized that, in children with a developing intestinal microbiome, evaluating microbiome composition in relation to chronological age might capture clinically relevant features not apparent from conventional measures of α-diversity or overall microbiome composition.

To define normal age-associated patterns of microbiome composition, we leveraged publicly available datasets comprising 1,313 fecal samples from healthy individuals spanning infancy through adulthood (**Fig. 4a**; **Suppl. Table 9**).^60–64^ We restricted this analysis to samples profiled by shotgun metagenomic sequencing to minimize technical variation across cohorts, given the substantial heterogeneity in 16S rRNA targeted regions, amplification protocols, and bioinformatic processing pipelines. Within the healthy cohort, chronological age was significantly associated with microbiome composition (PERMANOVA, *p*=0.001, **Fig. 4b**). Overall microbiome composition differed significantly between patients with HR-NB and healthy controls (Bray-Curtis PERMANOVA, *p*=0.001), although substantial overlap between the groups was observed in ordination space (**Fig. 4c**). Consistent with the association between age and microbiome composition, PCoA axis 1 was strongly correlated with chronological age in healthy children (Spearman, R=-0.64, *p*<0.001), whereas this association was weaker and did not reach statistical significance in pediatric HR-NB patients (R=-0.19, *p*=0.064; **Fig. 4d**).

**Figure 4.**
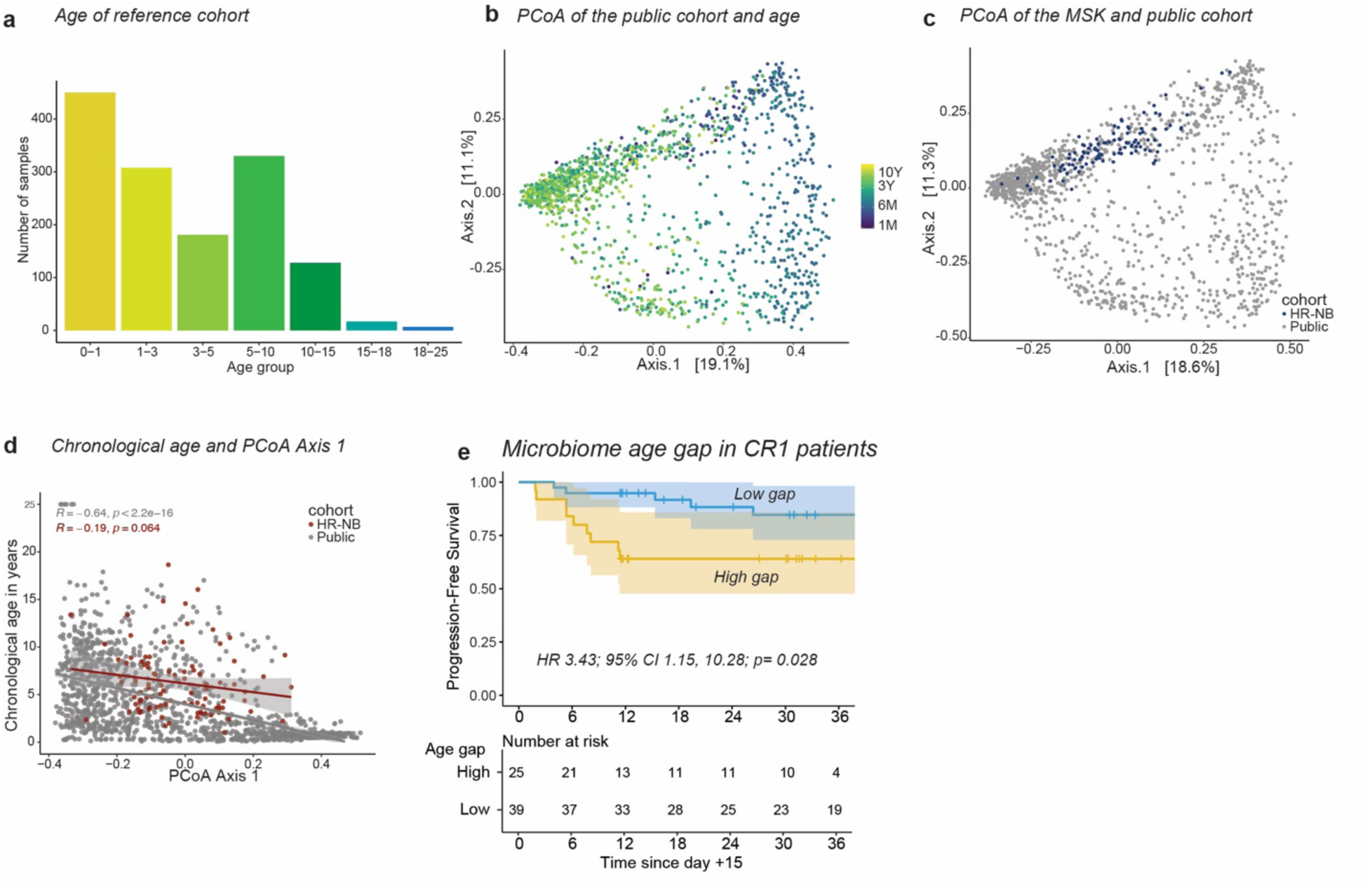
Increased microbiome relative age gap is associated with inferior progression-free survival in CR1. (**a**) Age dispersion in the public healthy cohort. (**b**) PCoA of the healthy cohort colored by age. (**c**) PCoA including all the healthy subjects and patients with HR-NB. (**d**) Correlation of chronological age with PCoA Axis 1. (**e**) Kaplan-Meier analysis of PFS according to microbiome relative age gap, dichotomized at the median, among patients in CR1. Hazard ratio and *p* value are derived from Cox regression. Statistical comparison in **d** was performed using Spearman correlation.

We hypothesized that the microbiome of patients with HR-NB would exhibit delayed maturation and therefore more closely resemble the microbiome of younger healthy children. For each patient, we identified three healthy reference samples with the most similar microbiome composition based on Bray-Curtis dissimilarity matrix and compared their chronological ages with that of the corresponding patient. In both CR1 and ≥CR2, microbiome-matched healthy children were significantly younger than the corresponding patients with HR-NB (median age difference 15.4 months in CR1, *p*<0.001; 44.3 months in ≥CR2, *p*<0.001; **Extended Data Fig. 4a,b**). To assess whether microbiome similarity recapitulated chronological age within the healthy cohort, we repeated the matching procedure using healthy subjects only. We randomly selected 94 healthy children and, for each child, identified the three healthy subjects with the most similar microbiome composition based on Bray-Curtis dissimilarity. The chronological age of the 94 selected children did not differ from the mean age of their microbiome-matched healthy subjects (*p*=0.64; **Extended Data Fig. 4c**), indicating that microbiome-based matching recapitulated age similarity within the healthy cohort. Together, these findings support that the microbiome composition of patients with HR-NB more closely resembled that of chronologically younger healthy children, consistent with delayed microbiome maturation.

We next quantified this age-related difference by calculating a microbiome relative age gap as the difference between each patient’s chronological age and the mean chronological age of their microbiome-matched reference samples, normalized to the patient’s chronological age. Values near zero indicate a microbiome resembling that of healthy children of similar chronological age, whereas increasingly positive values indicate resemblance to progressively younger healthy children and therefore greater maturational delay. Patients in ≥CR2 had significantly higher microbiome relative age gaps than patients in CR1 (*p*=0.002; **Extended Data Fig. 4d**), indicating greater microbiome maturational delay in ≥CR2 patients.

We then investigated whether microbiome maturation delay was associated with vaccine responses and clinical outcomes. Among CR1 patients, an above-median microbiome relative age gap was associated with inferior PFS (HR 3.43, 95% CI 1.15-10.28, *p*=0.028; **Fig. 4e; Suppl. Table 10**). This association was not observed in ≥CR2 patients (*p*=0.73, **Suppl. Table 10**). Microbiome relative age gap was also not associated with anti-GD2 IgG1 titers in either CR1 or ≥CR2 (Spearman, R=-0.01, *p*=0.96; R=-0.019, *p*=0.92, respectively; **Extended Data Fig. 4e**).

### Antibiotic-mediated microbiome perturbation impairs responses to the GD2/GD3 vaccine in mice

To investigate whether intestinal microbiome perturbation directly influences responses to GD2/GD3 vaccination, we performed preclinical experiments designed to recapitulate key features of the clinical study conditions.^65^ Wild-type C57BL/6 mice, 6-8 weeks of age, received subcutaneous GD2/GD3 vaccination with QS-21, using the same vaccine formulation administered to patients, followed by three booster vaccinations of the same formulation over 42 days. Mice also received β-glucan by oral gavage according to the treatment schedule shown in **Fig. 5a**. To perturb the intestinal microbiome, a subset of mice received vancomycin in their drinking water (0.5 g/L), selected for its activity against Gram-positive intestinal bacteria, including *Clostridia-*associated taxa.^66^

**Figure 5.**
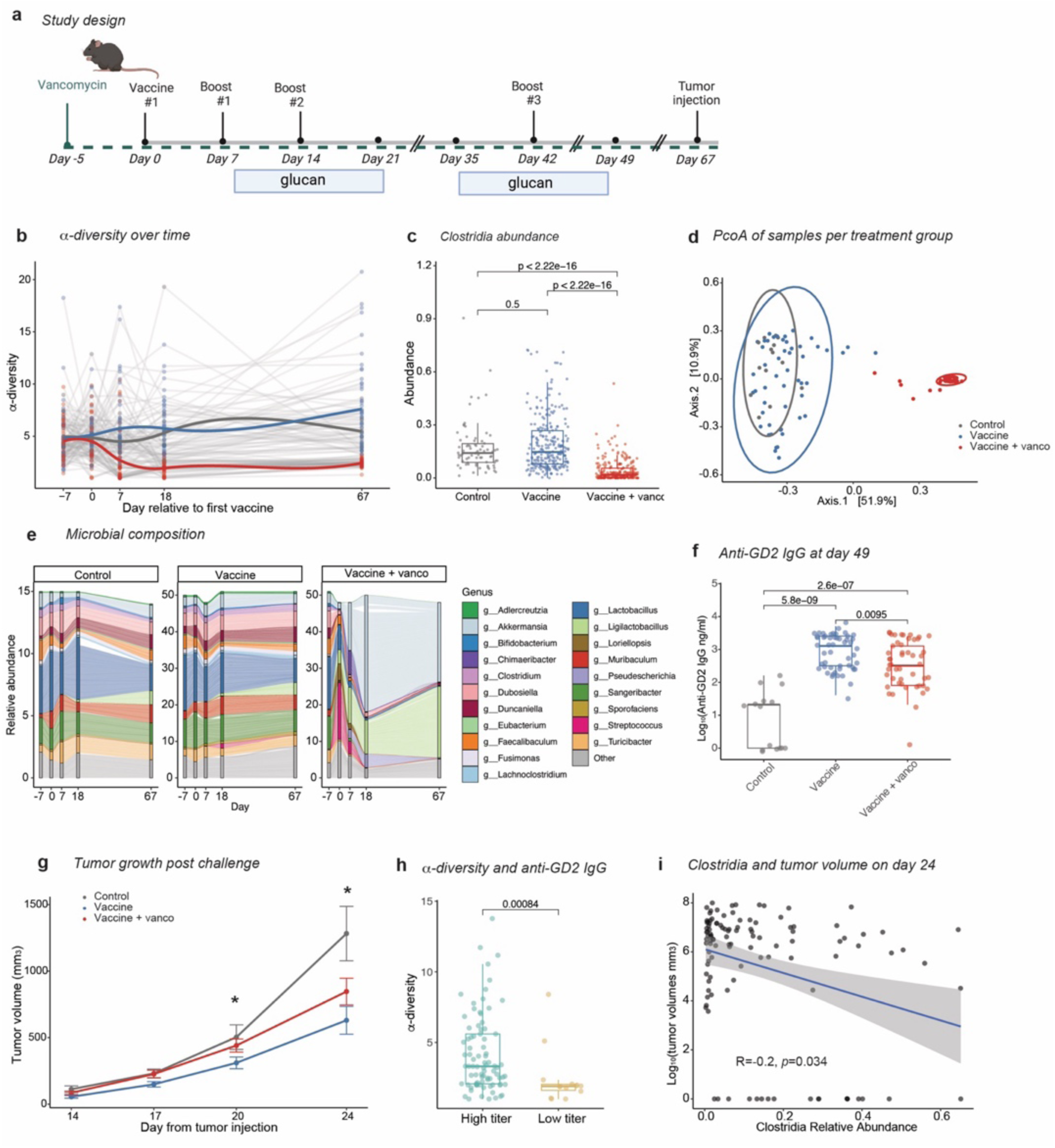
Antibiotic-mediated microbiome perturbation impairs vaccine-induced antibody responses and tumor control in mice. (**a**) Experimental schema of GD2/GD3 vaccination in wild type C57BL/6 mice. The green dotted line indicates continuous vancomycin administration in the drinking water. (**b**) Longitudinal α-diversity across treatment groups. (**c**) *Clostridia* abundance on day 18 according to treatment group. (**d**) PCoA of microbial community composition on day 18. (**e**) Microbial composition per treatment group over time. (**f**) Anti-GD2 IgG titers according to treatment group. (**g**) Tumor volume following tumor inoculation across treatment groups. (**h**) α-diversity and anti-GD2 IgG titers. Titers were dichotomized as positive or negative using a threshold derived from anti-GD2 IgG responses in control mice. (**i**) Association between *Clostridia* abundance at the time of tumor inoculation and tumor volume 24 days after inoculation. Data are pooled across five independent experiments and include 15 control mice, 51 vaccinated mice and 51 mice receiving vaccination plus vancomycin. Statistical comparisons in **c,f,h** were performed using a two-sided Wilcoxon rank-sum test, and in **i** were performed using Spearman correlation. Schematic (**a**) *created in BioRender. Miltiadous, O. (2026)* https://BioRender.com/cgg2n3u.

To assess the effects of vancomycin on the gut microbiome, we evaluated α-diversity in longitudinal stool samples. Baseline α-diversity was similar across treatment groups; however, within two weeks of antibiotic initiation, vancomycin-treated mice exhibited reduced α-diversity compared with control mice and mice receiving vaccine alone, and this reduction persisted throughout the experiment (**Fig. 5b, Extended Data Fig. 5a**). Vancomycin treatment also reduced the relative abundance of *Clostridia* (**Fig. 5c**). Overall microbial community composition differed following vancomycin treatment compared with control mice and mice receiving vaccine alone (PERMANOVA, *p*=0.001, **Fig. 5d,e**).

Vancomycin treatment impaired vaccine-induced humoral responses, with lower anti-GD2 IgG titers in mice receiving vancomycin plus vaccine compared with mice receiving vaccine alone (*p*=0.01, **Fig. 5f**). Following completion of the vaccine series, mice were challenged with 50,000 GD2-positive EL4 cells and subsequent tumor growth was monitored. Vancomycin-treated mice exhibited more rapid tumor growth than mice receiving vaccine alone, with significant differences observed on days 20 and 24 (**Fig. 5g, Extended Data Fig. 5b,c**). Consistent with our clinical findings, mice with positive anti-GD2 IgG responses had higher microbial α-diversity than mice with negative responses (*p*<0.001; **Fig. 5h**). Because antibody titers were measured as ordinal dilutional values, we also analyzed the association between α-diversity and anti-GD2 IgG responses using cumulative logistic regression. Higher α-diversity was associated with increasing anti-GD2 IgG titers (estimate=0.20, *p*=0.006). We next investigated whether the degree of relative *Clostridia* depletion was associated with tumor growth. We found a significant negative correlation between *Clostridia* abundance and tumor volume on day 24 (R=-0.2, *p*=0.034; **Fig. 5i**), consistent with the association between higher *Clostridia* abundance and superior PFS in patients with HR-NB.

Because mice originated from different vendor production sites and breeding rooms across experiments, we investigated whether these factors contributed to inter-experimental variability. Microbiome composition differed according to vendor production site and breeding room in ordination analysis, with significant differences across breeding rooms (PERMANOVA, *p*=0.001, **Extended Data Fig. 5d,e**). Anti-GD2 IgG titers and tumor responses also varied across experimental cohorts; however, the direction of the vancomycin treatment effect was consistent across experiments despite this inter-cohort heterogeneity (**Extended Data Fig. 5f,g**).

To investigate potential immunologic mechanisms underlying the impaired vaccine responses following vancomycin treatment, we performed flow cytometry of splenic and vaccine-draining lymph node immune populations at the end of the experiment, following tumor inoculation.

Compared with mice receiving vaccine alone, mice receiving vaccine plus vancomycin had reduced frequencies of splenic dendritic cells, particularly cDC1, together with a reduced frequency of CD19+ B cells in vaccine-draining lymph nodes (**Extended Data Fig. 6a,b**). These findings demonstrate that vancomycin-induced intestinal microbiome perturbation impaired humoral responses to GD2/GD3 vaccination and accelerated tumor growth, accompanied by alterations in immune populations implicated in vaccine responses.

## Discussion

In this study, we identify the intestinal microbiome as a potential modulator of therapeutic cancer vaccine responses in children with HR-NB. The GD2/GD3 vaccine combines ganglioside tumor antigens (GD2 and GD3) with the immunogenic carrier KLH and the saponin adjuvant QS-21, and is administered with oral β-glucan, an immunomodulatory polysaccharide.^12–18,67,68^ In the clinical cohort, microbiome-disrupting antibiotic exposure was associated with inferior PFS, while antibiotic-mediated microbiome perturbation impaired vaccine-induced antibody responses and accelerated tumor growth in mice. Among patients in CR1, enrichment of *Lachnospiraceae*, *Blautia*, and *Clostridia* was associated with higher anti-GD2 IgG1 responses or superior PFS, while enrichment of an *Erysipelatoclostridium* signature was associated with worse PFS. Children with HR-NB also exhibited microbiome profiles resembling those of chronologically younger healthy children, consistent with delayed microbiome maturation. Greater maturation delay was significantly associated with inferior PFS among patients in CR1. Together, these findings support the intestinal microbiome as a potentially modifiable factor influencing therapeutic cancer vaccine efficacy.

Our findings are consistent with observations from adult cancer immunotherapy studies, in which gut microbial composition has been associated with response to immune checkpoint blockade. In particular, *Lachnospiraceae, Blautia,* and *Clostridia* have been associated with favorable responses to PD-1-directed therapy and CAR T cell therapy, despite substantial heterogeneity in taxonomic signatures across studies.^26–29,69–71^ Conversely, antibiotic exposure has been associated with worse outcomes among patients receiving immune checkpoint blockade and CAR-T cell therapy.^26,28–30,72^ Several of these microbial taxa have also been associated with enhanced vaccine immunogenicity in preclinical models and clinical correlative studies,^73,74^ suggesting that they may support immune responses across different immunotherapies. We also found that a microbial signature characterized by *Erysipelatoclostridium* was associated with worse PFS among patients in CR1. Consistent with this finding, *Erysipelatoclostridium* has been associated with intestinal dysbiosis and unfavorable outcomes following cancer immunotherapy.^75–77^

Several mechanisms may explain these observations. Taxa associated with higher anti-GD2 IgG1 responses in this cohort have previously been linked to short-chain fatty acid production, suggesting a potential pathway through which these microbial communities could influence vaccine immunogenicity.^78–83^ Short-chain fatty acids can modulate multiple components of adaptive immunity, including dendritic cell activity and antigen presentation, B-cell responses and plasma cell differentiation, as well as T follicular helper cell function.^84–87^ Notably, both patients and mice received β-glucan, an immunomodulatory polysaccharide that can serve as a substrate for microbial fermentation and short-chain fatty acid production.^12,67^ β-glucan has been shown to promote antigen presentation and enhance antibody responses in preclinical vaccine models.^13–18,68^ Dendritic cells are central to antigen presentation and the initiation of adaptive immune responses, with cDC1s playing a central role in tumor-antigen cross-presentation and antitumor immunity.^88,89^ Whether these immunologic effects are mediated directly through the effects of β-glucan on immune cells, indirectly through short-chain fatty acids or other microbiome-derived metabolites, or through a combination of these mechanisms remains unclear.

Our preclinical findings provide experimental evidence that the intestinal microbiome influences responses to GD2/GD3 vaccination. Vancomycin-mediated microbiome perturbation depleted *Clostridia*-associated taxa, which were associated with favorable vaccine responses and clinical outcomes in patients. Vancomycin-treated mice had reduced anti-GD2 IgG responses and higher tumor burden after inoculation, together with reduced splenic dendritic cells, particularly cDC1, and reduced CD19+ B cells in vaccine-draining lymph nodes. In the context of carbohydrate-peptide conjugate vaccines, both B cells and dendritic cells have been shown to play a central role in antigen presentation to T cells,^90^ raising the possibility that similar mechanisms contribute to immune responses to the GD2/GD3 vaccine.

An important finding of this study is the association between microbiome maturation and clinical outcomes. Unlike adults, children have a dynamically developing intestinal microbiome, and intensive multimodal cancer therapy occurs during a critical period of microbial development. Children completing therapy for HR-NB exhibited microbiome compositions that more closely resembled those of younger healthy children, consistent with delayed microbiome maturation, and greater microbiome relative age gap was associated with inferior PFS among CR1 patients. These findings suggest that microbiome maturation may provide a pediatric-specific measure of microbiome health, capturing alterations in age-related microbial development that are not reflected by conventional measures such as α-diversity or individual taxonomic abundance.

Most associations between microbiome features and vaccine responses were observed among patients treated in CR1. The ≥CR2 cohort was smaller and more clinically heterogeneous, with multiple prior relapses and therapeutic exposures, potentially limiting our ability to detect consistent microbial associations. Differences between the cohorts may also reflect underlying disease biology, as aggressive tumor-intrinsic features in relapsed neuroblastoma may diminish the relative contribution of microbiome-mediated immune modulation to disease control. Notably, ≥CR2 patients exhibited substantially greater microbiome maturational delay than CR1 patients. This may reflect cumulative treatment-related microbial injury and a shift toward a more uniformly disrupted microbiome state, which could reduce our ability to detect associations with PFS within the ≥CR2 cohort. Together, these findings suggest that the influence and prognostic significance of the microbiome may vary according to disease state, with stronger associations observed among patients treated in first remission.

These findings have potential clinical implications; however, the mechanisms linking the microbiome to GD2/GD3 vaccine immunogenicity remain to be defined. We are currently investigating the role of microbial metabolites in shaping vaccine responses, with the goal of identifying microbiome-based interventions that could be evaluated in a clinical trial to enhance vaccine immunogenicity. Despite this mechanistic uncertainty, our findings suggest that antibiotic stewardship may be particularly important during therapeutic vaccination and that microbiome features may ultimately contribute to biomarkers of vaccine responsiveness.

Importantly, the association between the intestinal microbiome and clinical outcomes may reflect both effects on vaccine immunogenicity and effects on disease control that are independent of the measured vaccine-induced humoral response. This possibility is supported by the distinct patterns of association observed across endpoints: microbiome relative age gap was associated with PFS but not with anti-GD2 IgG1 titers, whereas α-diversity was associated with anti-GD2 IgG1 titers but not with PFS. We therefore hypothesize that the intestinal microbiome may influence outcomes through complementary pathways involving vaccine immunogenicity as well as mechanisms not captured by the measured humoral response. Further studies are needed to define the mechanisms underlying these associations.

More than a century ago, William Coley demonstrated that exposure to bacterial products could provoke antitumor immune responses, providing one of the earliest foundations for cancer immunotherapy.^91,92^ Although our understanding of the microbial signals that shape antitumor immunity remains incomplete, our findings suggest that the endogenous intestinal microbiome of children with cancer may influence the efficacy of therapeutic cancer vaccination. Together, the clinical and preclinical evidence presented here identifies the intestinal microbiome as a factor shaping therapeutic cancer vaccine responses, identifies microbiome maturation as a clinically relevant feature in pediatrics, and provides a rationale for investigating microbiome-directed strategies to improve vaccine immunogenicity.

## Methods

### Patient selection and clinical information

All patients enrolled in the GD2/GD3 vaccine study at Memorial Sloan Kettering Cancer center between 2017 and 2021 were approached for participation in longitudinal microbiome sampling. The study and biospecimen collection were approved by the Memorial Sloan Kettering Institutional Review Board under the protocols IRB 05-075, IRB 12-245, and IRB 16-834. Informed consent was obtained for all participants from a parent or legal guardian (or directly from participants who had reached the age of legal consent), with participant assent obtained when appropriate. Of 158 patients who consented to stool collection, 139 vaccine-course observations had at least one stool specimen available. Two patients contributed samples during two separate vaccine courses; to restrict analyses to one observation per patient, their subsequent vaccine courses were excluded, resulting in 137 unique patients included in the cohort. Patients received one primary GD2/GD3 vaccination followed by six booster vaccinations over one year, in combination with oral β-glucan administered on a two-weeks-on, two-weeks-off schedule (**Extended Data Fig. 2a**).^11^ Patient enrollment and clinical trial cohorts are summarized in **Fig. 1a, Suppl. Table 1** and **Extended Data Fig 1a,** with clinical characteristics presented in **Table 1** and **Suppl. Table 2**.

### Preclinical models and experimental design

Female C57BL/6 mice, aged 6-8 weeks, were obtained from The Jackson Laboratory production facilities in California or Maine and housed three mice per cage under standard conditions, with ad libitum access to food and water and a 12-h light–dark cycle. Experiments were conducted under an institutionally approved animal protocol. Three experimental groups were compared: control, vaccine alone, and vaccine plus vancomycin. Control mice did not receive the GD2/GD3 vaccine but received all other study interventions. Vancomycin (GoldBio, V-200-5) was administered in drinking water at a final concentration of 0.5 g/L. Drinking bottles were replaced weekly. Yeast β-glucan (Biotec Pharmacon, 210110-2) was administered by oral gavage at 40 mg/kg beginning on day 7 after the first vaccination. Mice received β-glucan five days per week for two weeks, followed by two weeks without treatment and a second two-week treatment course. Mice were vaccinated subcutaneously on days 0, 7, 14 and 42 with 5 μg GD2L/GD3L-KLH^65,93^ and 5 μg QS-21. Following completion of the vaccination series, mice were challenged with GD2-expressing EL4 tumor cells. EL4 cells were confirmed to express GD2 before *in vitro* expansion (99% GD2 positivity) and cultured in RPMI-1640 supplemented with 10% FBS and penicillin-streptomycin. On the day of tumor challenge, cells were collected, resuspended in PBS and mixed 1:1 with Matrigel (Corning, 356234). Each mouse received 50,000 EL4 cells in a total volume of 100 μl by subcutaneous injection into the right flank. Tumors were measured twice weekly using digital calipers, and tumor volume was calculated as length × width² × 0.5.

### Stool sample collection and processing

Patients: The majority of the samples were collected in the outpatient setting. Parents collected the samples using inverted toilet “hats” and transferred the stool into a container, secured in a biohazard zip-lock bag to prevent leakage, and stored in foam container equipped with pre-chilled freezer packs. These samples were either hand-delivered by the patient during a clinical appointment or transported from their home to the laboratory by courier service. For patients admitted to the hospital, stool samples were collected using inverted toilet “hats” and transferred into a container, which was then immediately delivered to the laboratory via a pneumatic tube system for refrigeration at 4°C. Upon receipt at the Molecular Microbiology Facility Laboratory at MSKCC, each specimen was assigned a unique identifier and logged into the fecal biobank database. Processing involved transferring approximately 0.5 mL of whole stool into four 2 mL cryovials using a clean, disposable spatula. These aliquots were then moved to a -80°C freezer for long-term storage until required for DNA extraction and downstream processing.

#### Mice

Fecal samples were collected at baseline (day -7) and on days 0, 7, 18 and 67 relative to the first vaccination. Samples were processed for microbiome profiling as described above with 16S rRNA sequencing.

#### Microbiome extraction and sequencing

For the extraction, microbial cells were lysed with silica bead-beating, followed by nucleic acids extraction. For 16S ribosomal-RNA gene sequencing, the V4-5 variable region was amplified and sequenced on the Illumina MiSeq platform.^94^ Amplicon sequence variants (ASVs) were inferred using the DADA2 pipeline^95^ and taxonomically assigned against the NCBI 16S rRNA sequence database using BLAST.^95–97^ Microbial α-diversity was quantified using the inverse Simpson index.

For shotgun metagenomic sequencing, libraries were prepared using the Illumina TruSeq DNA library preparation kit and sequenced on the Illumina platform, targeting 10-20 million reads per sample.^98^ Metagenomic sequencing data were preprocessed with https://github.com/vdblab-shotgun/ “preprocess” mode, version 0.4.0. In short, data were deduplicated with bbtools’ clumpify; bbduk was used for quality trimming and artifact removal, and host-depletion was performed against human and mouse reference genomes (chm13v2.0 and GRCm39, respectively) with both Bowtie2 and SNAP.^99^ Taxonomic profiling was performed with Metaphlan 4.0.5 (database version mpa_vJan21_CHOCOPhlAnSGB_202103) using https://github.com/vdblab-shotgun/ “biobakery” mode, version 0.4.2.

### Anti-GD2 ELISA

#### Patients

Anti-GD2 IgG1 titers were quantified 7 weeks from the start of the vaccine, by ELISA using GD2-coated microtiter plates as previously described.^11^ GD2 (20 ng per well) was placed on 96-well plates, followed by blocking with 0.5% bovine serum albumin (BSA) for 1 h at room temperature (RT). Diluted patient serum was then added and incubated for 2.5 h at 37°C. After washing, bound anti-GD2 IgG1 was detected with a peroxidase-conjugated mouse anti-human IgG1 antibody during a 1h incubation at 4°C. Absorbance was measured at 490 nm. Hu3F8-IgG1 served as the reference standard for quantification, and the assay had a sensitivity of 0.6 ng/mL. Anti-GD2 IgG1 titer data were available for 125 of 137 patients.

#### Mice

Serum was collected by retro-orbital bleeding on day 49 for measurement of anti-GD2 IgG titers. Blood was centrifuged at 15,000 rpm for 15 min to isolate serum and frozen at -20C. Polystyrene microtiter ELISA plates (96 well) were coated overnight at 4C with 0.2 μg/well GD2 capture antigen for detecting antibodies in the mouse sera. The coated wells were then blocked with 200 μl/well of 1% HSA for 2 hours at RT. Sera from mice were serially diluted into 96 well plates and incubated with the antigen coated plate for 1.5h at RT. After washing five times with 1X PBS-Tween20, 100 μl/well of alkaline phosphatase-labeled secondary antibody conjugate with goat anti-mouse IgG (Southern Biotech, 1030-04) was added and incubated for 1.5h at RT.

Alkaline phosphatase substrate (p-Nitrophenyl phosphate; Sigma N1891) was added to each well and the plates were incubated for 45 min at room temperature. Absorbance at 405 nm was measured in an ELISA plate reader. Serial dilutions and standard curves were performed to calculate antibody concentrations. Anti-GD2 monoclonal or GD2-positive sera samples were used as positive controls.

### Flow cytometry

Mice were euthanized 25 days after tumor inoculation, and spleens and vaccine-draining lymph nodes were harvested and processed into single-cell suspensions in RPMI-1640. Cells were incubated with Fc receptor-blocking reagent and subsequently stained with DAPI and antibodies against CD19, CD3e, B220, MHC-II, CD45.2, CD8α, CD11c, CD11b, NK1.1, CD4, F4/80, CD172a (SIRPα) and XCR1. Antibody clones, fluorophores, manufacturers and catalog numbers are provided in **Suppl. Table 11**. Flow cytometric data were acquired using a BD FACSymphony A5 and a Cytek Aurora spectral flow cytometer and analyzed using FlowJo version 11. Representative gating strategy shown in **Extended Data Fig. 6c**.

### Statistical analysis

Microbial community composition was evaluated at the genus level using PCoA, and differences in community composition between groups were assessed using permutational multivariate analysis of variance (PERMANOVA).^100^ Topic modeling was used to identify microbial community signatures comprising taxa that tended to co-occur across samples.^101^ Differentially abundant taxa between prespecified patient groups were identified with LEfSe.^59^ Prior to these analyses, taxa with a relative abundance <1% or a prevalence <10% were grouped into a rare-taxa category at the highest taxonomic rank. An FDR threshold of 0.05 was applied to account for multiple testing, and an LDA score ≥4 was required to identify differentially abundant taxa. To assess differential taxonomic abundance according to 12-month relapse status, patients were classified as relapsed if relapse occurred within 12 months of vaccine initiation and as in remission if they remained relapse-free at 12 months. Patients with less than 12 months of follow-up who had not experienced a relapse were excluded from this analysis.

Continuous variables were compared between groups using two-sided Wilcoxon rank-sum tests, and *p*<0.05 was considered statistically significant. Associations between continuous variables were assessed using Spearman correlation. Linear regression was performed to evaluate factors associated with anti-GD2 IgG1 titers. Because anti-GD2 IgG titers in mice were measured by serial dilution and analyzed as ordinal values, associations with antibody responses were assessed using either a dichotomized titer threshold or cumulative logistic regression. The threshold for a positive anti-GD2 IgG response was defined as the mean titer plus two standard deviations in control mice. Multiple-testing correction was applied where appropriate, and results were considered statistically significant at an adjusted *p* value <0.05. Progression-free survival was estimated using the Kaplan-Meier method, and associations between microbiome features and PFS were evaluated using Cox proportional-hazards regression. Variables with *p*<0.20 in univariable analyses were included in the corresponding multivariable Cox models.

An independent publicly available cohort of healthy children and young adults spanning the age range of the MSK cohort was used as a reference for microbiome age (**Suppl. Table 9**). For each patient with HR-NB, the three healthy reference samples with the most similar microbiome composition were identified based on Bray-Curtis distance. Healthy reference samples were matched to HR-NB patients, such that the same healthy individual could serve as a microbiome match for more than one patient, thereby preserving the closest available microbiome matches across patients without restricting matches based on prior assignment. Matching was based solely on microbiome similarity and did not incorporate chronological age.

The microbiome relative age gap was calculated as the difference between each patient’s chronological age and the mean chronological age of their three microbiome-matched healthy controls, normalized to the patient’s chronological age: (patient age – mean age of microbiome-matched healthy controls)/patient age. Positive values indicate a microbiome resembling that of younger healthy individuals, with increasingly positive values indicating greater microbiome maturational delay; values near zero indicate greater resemblance to healthy individuals of similar chronological age.

## Acknowledgments

We thank Audrey Mauguen for biostatistical consultation. We thank the Molecular Microbiology Facility, including Emily Fontana, Luigi Amoretti, Ana Grandissimo, Ariel Molina, Jennifer Haber, Minsu Jaun, for their support with DNA extraction from stool samples; the Integrated Genomics Operation (iGO) for sequencing patient samples; and Novogene for sequencing murine stool samples.

## Author Contributions

L.M. performed computational analyses and contributed to manuscript preparation. K.A. and M.B.S. contributed to the design of the preclinical experiments and analysis of flow cytometry data. K.D., X.W., Y.Z., L.B., K.L., C.N., K.V., J.Y., and G.R. assisted with preclinical experiments. N.R.W. and A.D. provided computational support. T.F. provided biostatistical support and oversight. I.Y.C. provided anti-GD2 IgG1 titer data from patient samples. L.H. and N.B. assisted with patient enrollment and coordinated fecal sample collection. J.U.P., K.A.M., Y.T., J.J.B., S.N., G.D., I.M., N.K.C., M.V.B., and S.M. contributed to analytical strategies and interpretation of the findings. S.S.R., B.H.K. and S.M. contributed to clinical protocol design and patient enrollment. O.M. conceptualized and supervised the study, contributed to data analysis and interpretation, and wrote the manuscript. All authors reviewed and approved the final manuscript.

## Data availability

The patient and murine microbiome sequencing data generated in this study are deposited in the NCBI Sequence Read Archive (SRA) under the BioProject PRJNA1537701 and will be publicly available upon publication. Publicly available metagenomic datasets used to construct the healthy reference cohort are listed with their corresponding accession numbers in **Suppl. Table 9**. Additional data and code supporting the findings of this study will be made available upon publication.

