## Extended Data Figures for "The intestinal microbiome shapes responses to the GD2/GD3 cancer vaccine in children with high-risk neuroblastoma"

**Extended Data Figure 1. Study design, longitudinal sample collection and clinical trial primary endpoints.** (**a**) CONSORT diagram showing patients enrolled in the Phase II and Phase II randomization cohorts and their overlap with the microbiome study population. (**b-e**) Clinical outcomes according to trial cohort. (**b**) One-year event-free survival among patients enrolled in the Phase II CR1 cohort. (**c**) Six-month clinical response status among patients enrolled in the Phase II ≥CR2 cohort. (**d**) Week 32 anti-GD2 IgG1 antibody titers among patients randomized to initiate β-glucan at week 1 versus week 6. The analysis included 55 patients with available week 32 titers, of whom n=31 initiated β-glucan at week 1 and n=24 at week 6. (**e**) Week 8 anti-GD2 antibody titers according to β-glucan randomization group. The analysis included 70 patients with available week 8 titers, of whom n=38 initiated β-glucan at week 1 and n=32 at week 6. Statistical comparisons in **d** and **e** were performed using linear regression adjusted for CR status; the week 32 comparison in **d** represents the prespecified primary analysis.

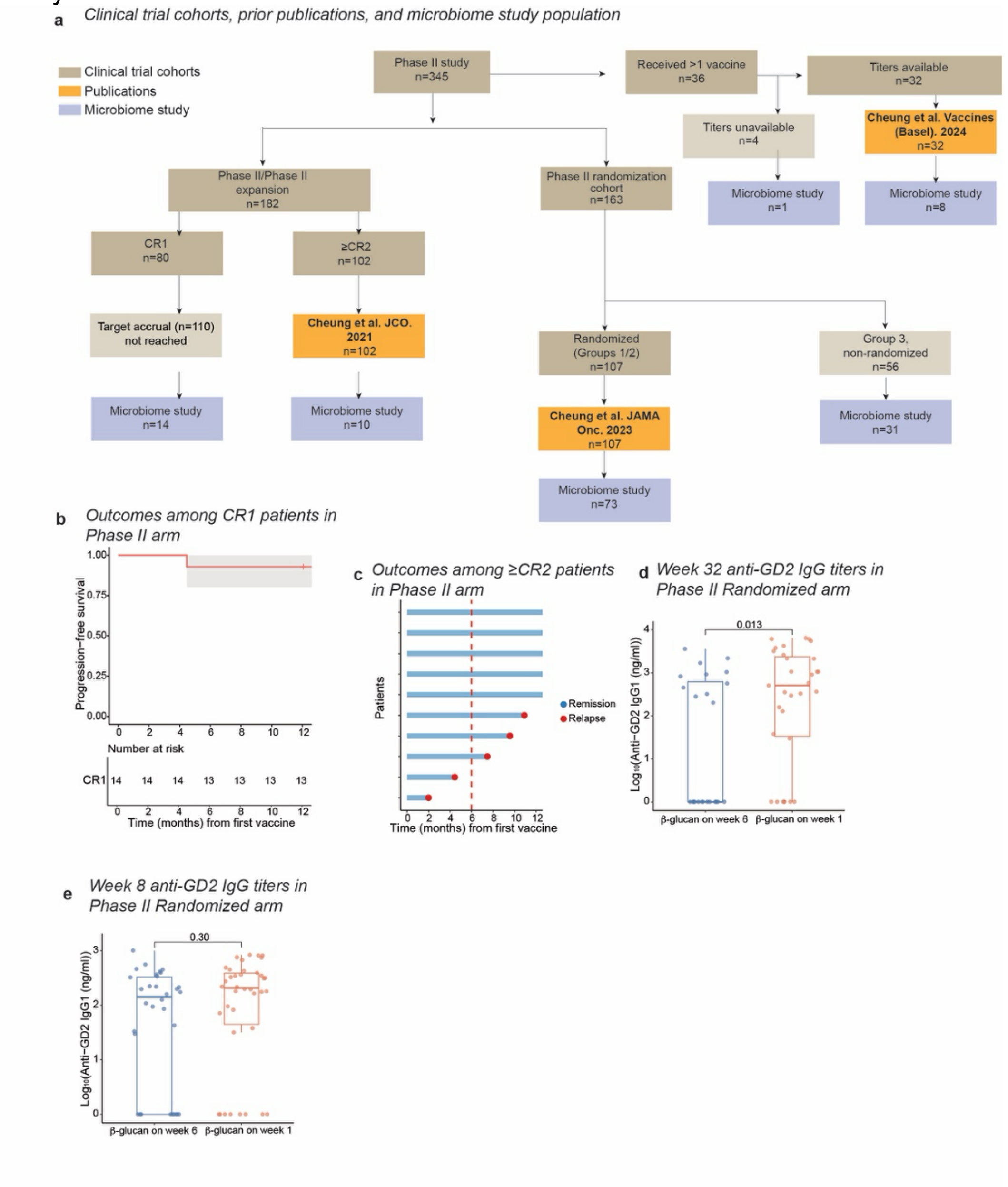


**Extended Data Fig 1.**

**Extended Data Figure 2. Antibiotic exposure and baseline microbiome composition in the clinical cohort.** (**a**) Study design. (**b**) Longitudinal stool sample collection throughout the study for the 137 patients enrolled in the microbiome cohort. Each dot represents an individual stool sample, with samples from the same patient connected by a line. Dashed vertical lines indicate the timing of the seven vaccine doses. (**c**) Paired comparison of α-diversity in stool samples from seven patients collected before and after exposure to moderate/high microbiome-perturbing antibiotics, with samples collected within 15 days of antibiotic administration. (**d**) Temporal relationship between antibiotic exposure and stool sample collection. (**e**) Schematic illustrating the relationship between the timing of antibiotic exposure and stool collection and the expected microbiome composition captured under each sampling scenario. (**f**) PCoA of baseline microbial community composition stratified by remission status (CR1 versus ≥CR2). Points are colored according to age (<5 versus ≥5 years), with point shape indicating prior hematopoietic cell transplantation (HCT). Statistical comparison in **c** was performed using a two-sided Wilcoxon signed-rank test.

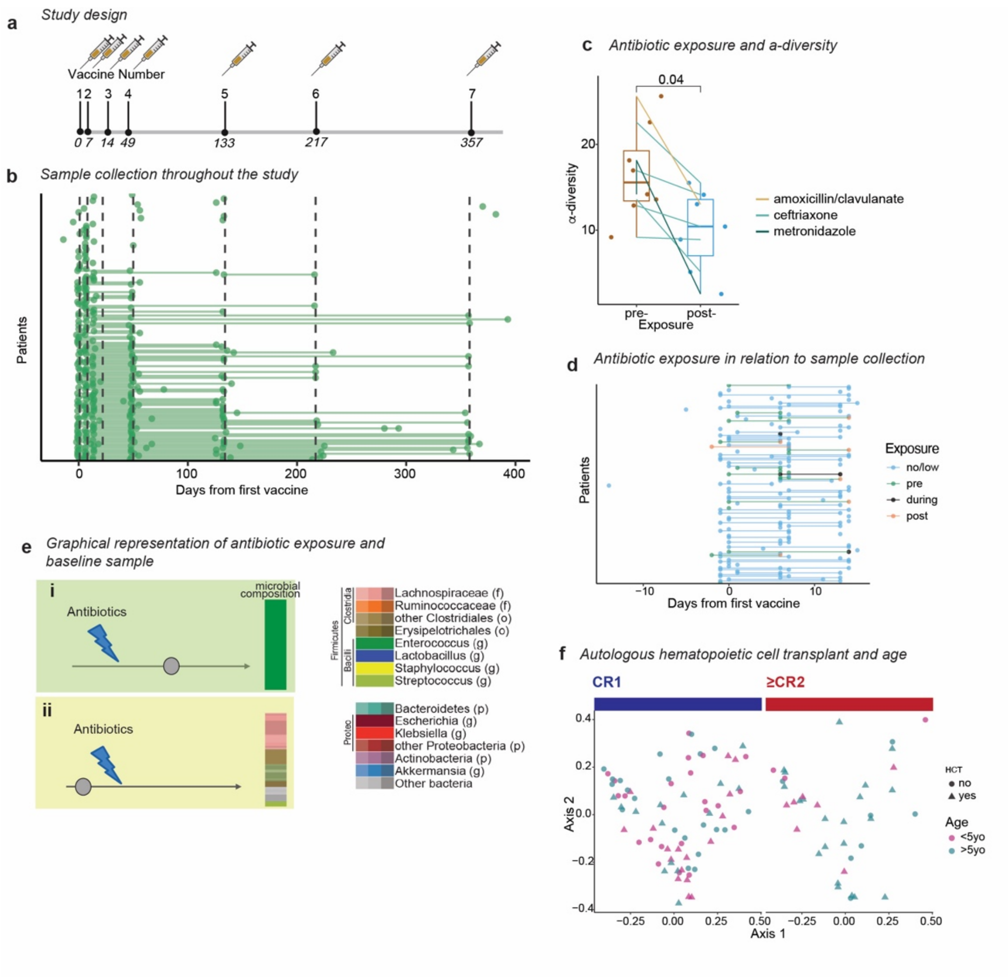


**Extended Data Fig 2.**

**Extended Data Figure 3. Microbial community signatures**. (**a**) Microbial community signatures identified by topic modeling and labeled according to the dominant taxon within each signature for ≥CR2. PCoA of baseline samples colored by the topic signature in (**b**) CR1 and (**c**) ≥CR2 patients. (**d**) Distribution of *Erysipelatoclostridium* signature among CR1 patients. (**e**) Temporal distribution of relapses following initiation of GD2/GD3 vaccination.

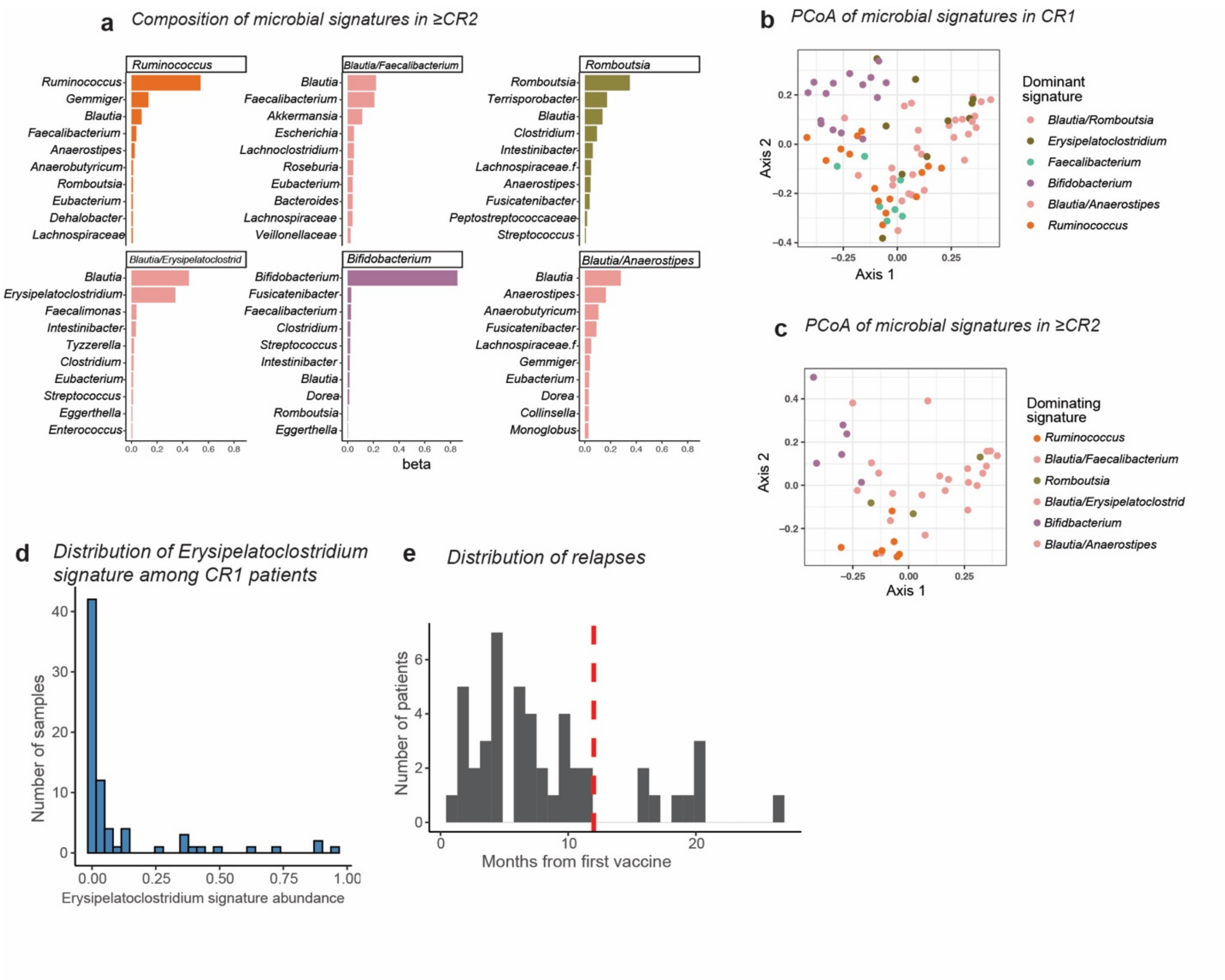


**Extended Data Fig. 3.**

**Extended Data Figure 4. Microbiome relative age gap**. (**a-b**) Comparison of chronological age between patients in CR1 and ≥CR2 and their microbiome-matched healthy controls. (**c**) Chronological age of healthy children and microbiome-matched healthy subjects. (**d**) Comparison of Microbiome relative age gap in patients vaccinated in CR1 vs ≥CR2. (**e**) Microbiome age gap and anti-GD2 IgG1 titers. Statistical comparison in **a-c** was performed using a two-sided Wilcoxon signed-rank test, **d** using a two-sided Wilcoxon rank-sum test, and in **e** using Spearman correlation.

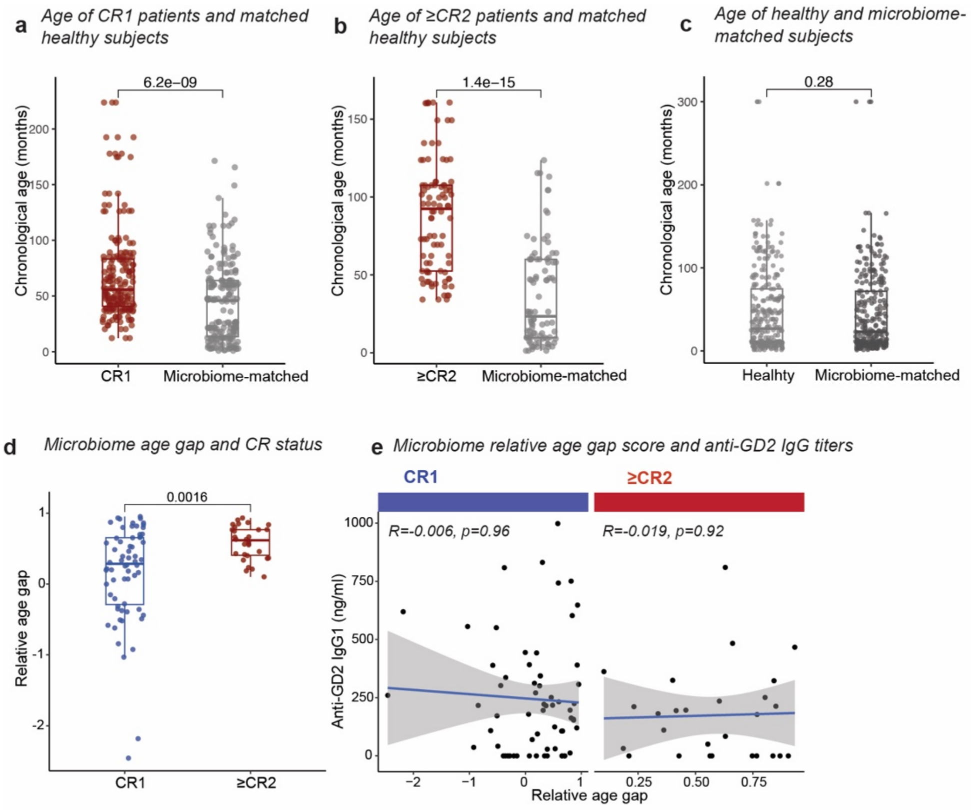


**Extended Data Fig 4.**

**Extended Data Figure 5. Microbiome and vaccine responses in preclinical models.** (**a**) α-diversity comparisons across treatment groups throughout the experiment. (**b**) Individual tumor growth trajectories stratified by treatment group. (**c**) Tumor growth across treatment groups from days 17 to 24 after tumor inoculation. (**d**) Microbial composition across experimental cohorts and treatment groups. (**e**) PCoA of microbial community composition with samples colored according to experimental cohort and production site/breeding room. (**f**) Tumor growth per treatment group and production site. (**g**) Anti-GD2 IgG titers stratified by treatment group and production site/breeding room. Statistical comparisons in **a,c,g** were performed using pairwise two-sided Wilcoxon rank-sum tests.

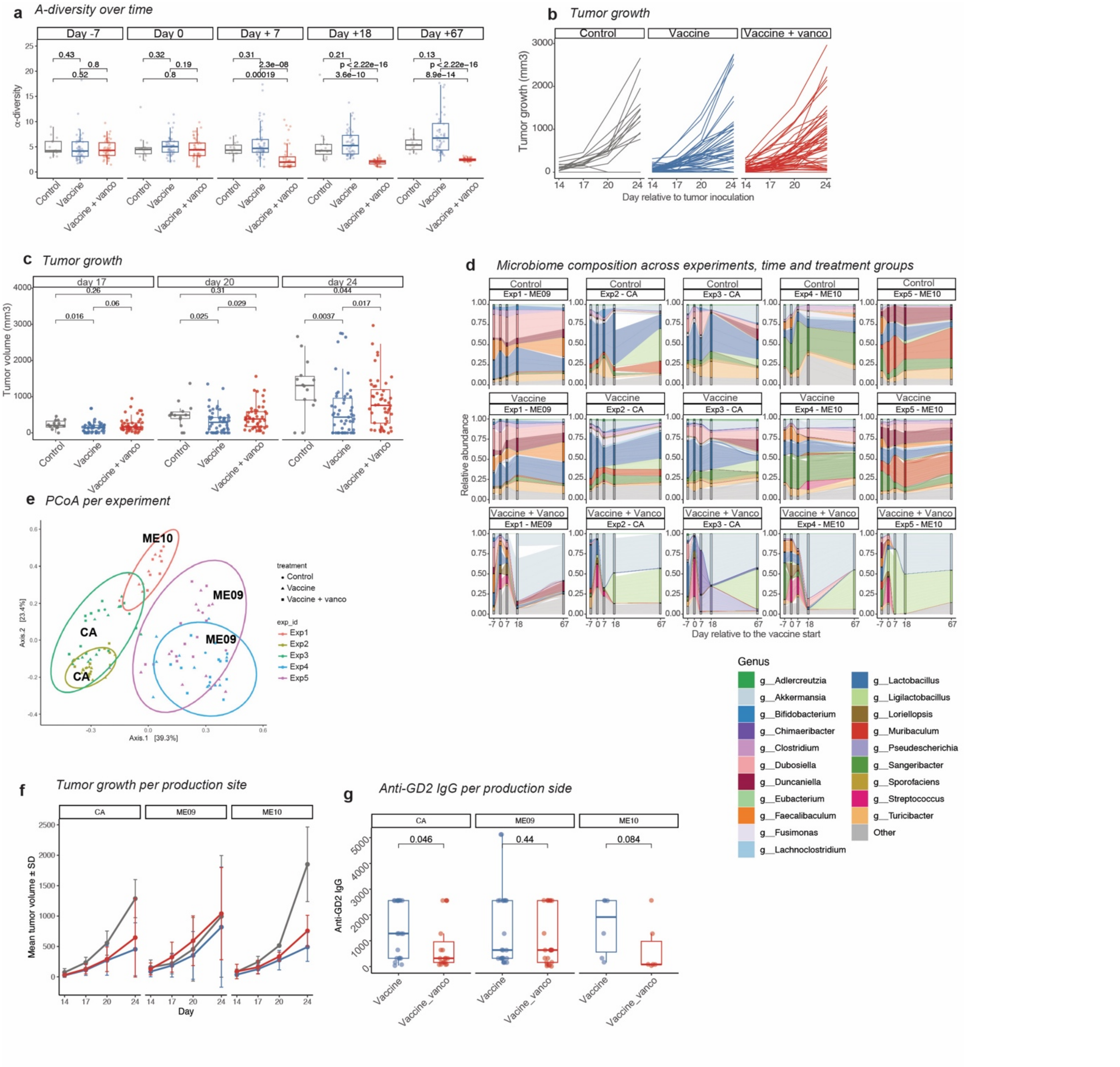


Extended Data Fig. 5

**Extended Data Figure 6. Immune cell populations following GD2/GD3 vaccination and tumor challenge.** (**a**) Absolute counts and frequencies of dendritic cells (DCs), conventional type 1 dendritic cells (cDC1) and conventional type 2 dendritic cells (cDC2) in the spleen across treatment groups. Frequencies are expressed as percentages of the respective parent immune cell population. (**b**) Absolute counts and frequencies of CD19+ B cells in vaccine-draining lymph nodes across treatment groups, with frequencies expressed as percentages of CD45+ cells. Immune populations were evaluated at the end of the experiment, 25 days after tumor inoculation, across two independent experiments, comprising 6 control mice, 24 vaccinated mice and 24 mice receiving vaccination plus vancomycin. Samples were excluded if no viable cells were available for flow cytometry. Statistical comparisons were performed using two-sided Wilcoxon rank-sum tests.

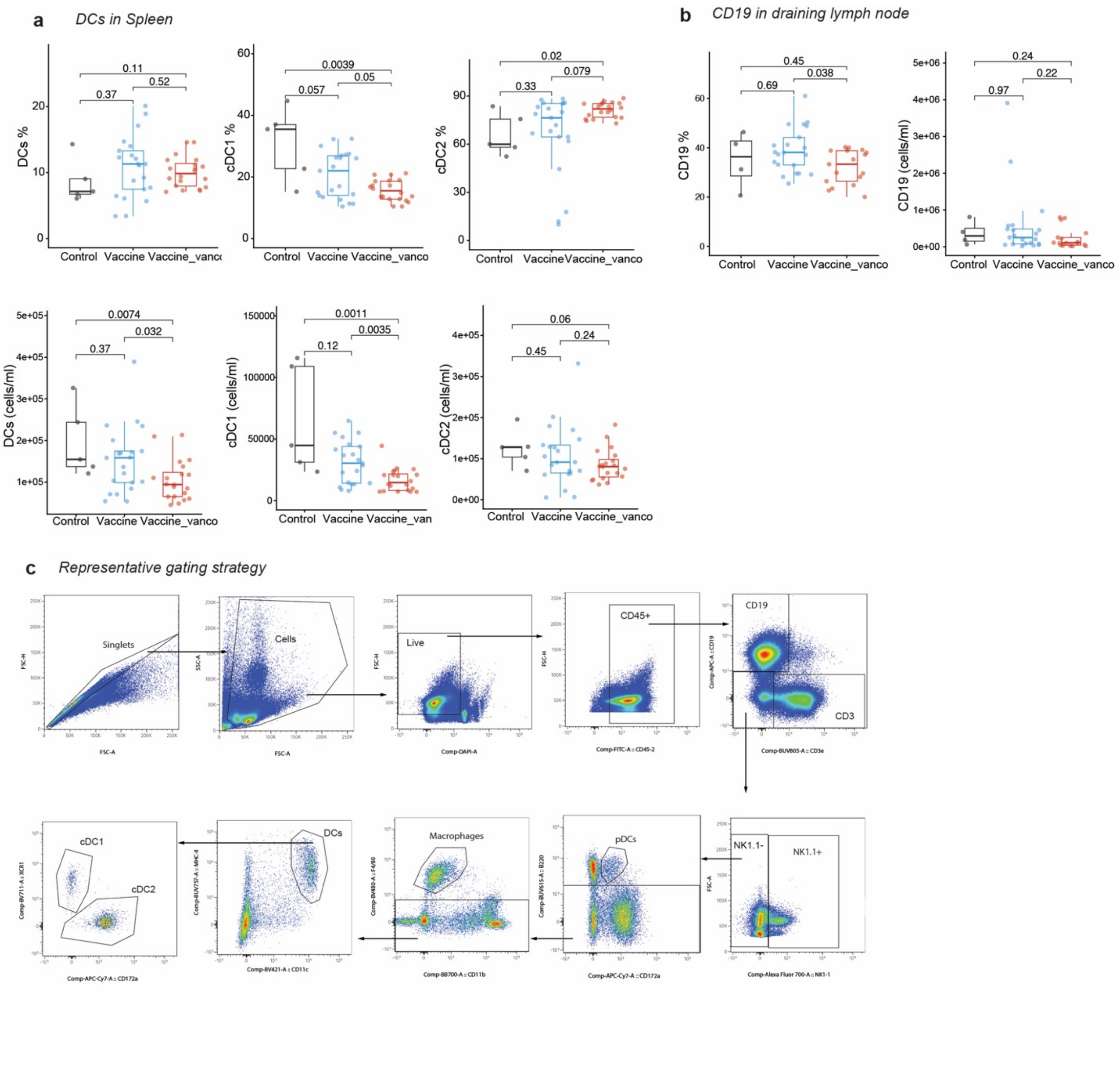


Extended Data Fig. 6
