## Supplementary Tables for "The intestinal microbiome shapes responses to the GD2/GD3 cancer vaccine in children with high-risk neuroblastoma"

**Supplementary Table 1.** Clinical trial cohorts and protocol-defined objectives relevant to patients represented in the microbiome study.

| Trial subcohort | Microbiome cohort (n) | Protocol-defined primary outcome | Previous reporting |
|---|---|---|---|
| **Phase II#** | | | |
| CR1 | n=14 | 1-year RFS | No; target accrual not reached |
| ≥CR2 | n=10 | 6-month CR | Yes; *Cheung et al., JCO 2021*[2] |
| **Phase II randomization cohort** | | | |
| Phase II Group 1/2, randomized | n=73 | Post-glucan 32-week anti-GD2 response, adjusted for CR status | Yes; *Cheung et al., JAMA 2023*[1] |
| Phase II Group 3, nonrandomized | n=31 | Exploratory cohort* | No |
| **>1 vaccine** | n=9 | Exploratory cohort* | Yes; *Cheung et al., Vaccines 2024*[3] |

#No minimal residual disease (MRD) positive patients were included in the microbiome study. Therefore, the protocol-defined primary objective assessing anti-neuroblastoma activity in patients with MRD was not represented in the microbiome cohort.
*Exploratory sub-cohorts without a separate protocol-defined primary outcome.
RFS, relapse-free survival; CR, complete remission.

**Supplementary Table 2.** Baseline clinical characteristics of the clinical trial subcohorts of patients included in the microbiome study.

| | Phase II | | Phase II Randomized | | Non-randomized, Group 3 | | Vaccine >1 | Total (n=137) |
|---|---|---|---|---|---|---|---|---|
| **Sub-cohort** | **CR1 (n=14)[1]** | **≥CR2 (n=10)[1]** | **CR1 (n=51)[1]** | **≥CR2 (n=22)[1]** | **CR1 (n=24)[1]** | **≥CR2 (n=7)[1]** | **≥CR2 (n=9)[1]** | |
| **Age at diagnosis (mo)** | 31 (29,50) | 40 (21,59) | 41 (25,64) | 46 (27,68) | 34 (27,57) | 24 (15,55) | 33 (24,81) | 36 (25, 60) |
| **Age at vaccine entry (mo)** | 47 (44,63) | 101 (62,110) | 61 (39,83) | 89 (52,108) | 62 (47,91) | 66 (56,110) | 91 (83,123) | 66 (46, 98) |
| **Sex (M)** | 5 (36%) | 10 (100%) | 28 (55%) | 13 (59%) | 13 (54%) | 5 (71%) | 2 (22%) | 76 (55%) |
| **Stage (M)** | 14 (100%) | 10 (100%) | 48 (94%) | 22 (100%) | 24 (100%) | 7 (100%) | 8 (89%) | 133 (97%) |
| ***MYCN* amplification** | 6 (43%) | 2 (20%) | 15 (32%) | 4 (19%) | 9 (41%) | 4 (57%) | 2 (25%) | 42 (33%) |
| *MYCN* unknown | 0 | 0 | 4 | 1 | 2 | 0 | 1 | 8 |
| **HCT** | 6 (43%) | 7 (70%) | 22 (43%) | 16 (73%) | 11 (46%) | 6 (86%) | 4 (44%) | 72 (53%) |
| **Number of previous relapses** | | | | | | | | |
| **0** | 14 (100%) | 0 (0%) | 51 (100%) | 0 (0%) | 24 (100%) | 0 (0%) | 0 (0%) | 89 (65%) |
| **1** | 0 (0%) | 6 (60%) | 0 (0%) | 21 (95%) | 0 (0%) | 6 (86%) | 6 (67%) | 39 (28%) |
| **2** | 0 (0%) | 1 (10%) | 0 (0%) | 1 (4.5%) | 0 (0%) | 1 (14%) | 2 (22%) | 5 (3.6%) |
| **3** | 0 (0%) | 3 (30%) | 0 (0%) | 0 (0%) | 0 (0%) | 0 (0%) | 1 (11%) | 4 (2.9%) |

**Supplementary Table 3.** Classification of antibiotics by microbiome-perturbing potential.

| | |
|---|---|
| **Low perturbation** | |
| | Levofloxacin |
| | Sulfamethoxazole-trimethoprim |
| **Moderate/High perturbation** | |
| | Amoxicillin |
| | Amoxicillin-Clavulanate |
| | Ceftriaxone |
| | Metronidazole |

**Supplementary Table 4.** Indications for antibiotic administration.

| UPN | Age (mo) | *MYCN* ampl | Auto-HCT | CR | Antibiotic | Reason | Infection | Start | End | Hospital status |
|---|---|---|---|---|---|---|---|---|---|---|
| 2748 | 28 | amp | no | CR1 | amoxicillin | strep throat | Yes | -20 | -11 | outpatient |
| 7432 | 190 | no | no | ≥CR2 | ceftriaxone | central line infection | Yes | -24 | -14 | inpatient |
| 2730 | 110 | no | yes | ≥CR2 | metronidazole | C. Diff | Yes | 6 | 15 | outpatient |
| 2751 | 42 | amp | yes | CR1 | amoxicillin/clavulanate | thumb fracture | No | 1 | 1 | outpatient |
| 2727 | 108 | no | yes | ≥CR2 | ceftriaxone | fever | No | 8 | 8 | outpatient |
| 7193 | 110 | no | no | ≥CR2 | ceftriaxone | fever | No | 1 | 1 | outpatient |
| 2707 | 72 | no | yes | CR1 | ceftriaxone | fever | No | 15 | 15 | outpatient |
| 2680 | 37 | no | yes | ≥CR2 | ceftriaxone | fever | No | 14 | 14 | outpatient |
| 2711 | 73 | no | yes | CR1 | ceftriaxone | fever | No | 6 | 6 | outpatient |
| 2661B | 72 | no | no | ≥CR2 | ceftriaxone | fever | No | 1 | 1 | outpatient |
| 2665 | 100 | no | yes | ≥CR2 | levofloxacin | fever | No | -2 | -2 | outpatient |
| 2764 | 88 | no | yes | ≥CR2 | ceftriaxone | fever | No | 8 | 8 | outpatient |
| 2746 | 35 | amp | yes | CR1 | ceftriaxone | fever | No | 8 | 8 | outpatient |
| 2731 | 74 | no | yes | CR1 | ceftriaxone | fever | No | 8 | 8 | outpatient |
| 2734 | 76 | no | yes | CR1 | ceftriaxone | fever | No | 8 | 8 | outpatient |
| 2728 | 53 | amp | yes | CR1 | ceftriaxone | fever | No | 8 | 8 | outpatient |
| 2769 | 106 | amp | no | CR1 | ceftriaxone | fever | No | 8 | 8 | outpatient |
| 2766 | 155 | no | yes | ≥CR2 | ceftriaxone | fever | No | 8 | 8 | outpatient |
| 7866 | 42 | amp | no | CR1 | amoxicillin | fever | No | 14 | 24 | outpatient |
| 2678B | 153 | no | no | ≥CR2 | ceftriaxone | fever | No | 1 | 1 | outpatient |
| 2756 | 86 | no | no | CR1 | ceftriaxone | fever | No | 6 | 6 | outpatient |
| 2743 | 66 | amp | yes | ≥CR2 | ceftriaxone | fever | No | 8 | 8 | outpatient |
| 2732 | 40 | amp | no | CR1 | ceftriaxone | fever | No | 8 | 8 | outpatient |

**Supplementary Table 5.** Cox regression analysis of PFS.

| | | *Univariable analysis* | | | | *Multivariable analysis* | |
|---|---|---|---|---|---|---|---|
| | **n** | **HR** | **95% CI** | ***p*-value** | **HR** | **95% CI** | ***p*-value** |
| **Males** (ref: females) | 137 | 1.26 | 0.70, 2.26 | 0.44 | | | |
| **Age at vaccine** (years) | 137 | 1.13 | 1.06, 1.21 | <0.001 | 1.10 | 1.00, 1.20 | 0.041 |
| ≥**CR2** (ref: CR1) | 137 | 4.00 | 2.22, 7.22 | <0.001 | 3.29 | 1.73, 6.27 | <0.001 |
| **HCT** (ref: no HCT) | 137 | 0.83 | 0.47, 1.48 | 0.54 | | | |
| **Glucan initiated day 1** (ref: day 35) | 137 | 0.99 | 0.53, 1.84 | 0.97 | | | |
| **MYCN amplified** (ref: non-amplified) | 129 | 0.60 | 0.29, 1.21 | 0.15 | 0.98 | 0.46, 2.13 | 0.968 |
| **Mod/high perturbation antibiotics** (ref: no/low perturbation antibiotics) | 137 | 2.11 | 1.09, 4.07 | 0.03 | 2.02 | 1.03, 3.98 | 0.041 |

*HRs were estimated using Cox proportional hazards regression models for progression-free survival. Reference categories for categorical variables are indicated in parentheses. Variables with p<0.20 in univariable analyses were included in the corresponding multivariable model. HR, hazard ratio; CI, confidence interval; CR, complete remission; HCT, hematopoietic cell transplantation.*

**Supplementary Table 6**. General linear model evaluating associations with anti-GD2 IgG1 titers.

| | Univariable **analysis** | | | | **Multivariable analysis** | | |
|---|---|---|---|---|---|---|---|
| **Characteristic** | n | β | 95% CI | p-val | β | 95% CI | p-val |
| **CR1** | | | | | | | |
| **Males** (ref: females) | 73 | 0.19 | -0.31, 0.69 | 0.44 | | | |
| **Age at vaccine** (years) | 73 | -0.001 | -0.07, 0.07 | 0.97 | | | |
| **Glucan initiated day 1** (ref: day 35) | 73 | 0.27 | -0.26, 0.81 | 0.31 | | | |
| ***HCT** (ref: no HCT)* | *73* | *-0.40* | *-0.90, 0.09* | *0.11* | *-0.51* | *-0.98, -0.04* | *0.03* |
| **MYCN amplified** (ref: non-amplified) | 67 | -0.56 | -1.08, -0.04 | 0.04 | -0.43 | -0.90, 0.05 | 0.08 |
| ***α-diversity*** | *73* | *0.04* | *-0.002, 0.09* | *0.06* | *0.07* | *0.03, 0.11* | *0.002* |
| **Signatures** | 73 | | | | | | |
| *Blautia/Romboutsia* | | 0.17 | -0.98, 1.33 | 0.77 | | | |
| *Erysipelatoclostridium* | | *-0.07* | *-1.18, 1.05* | *0.90* | | | |
| *Faecalibacterium* | | -0.83 | -2.27, 0.62 | 0.26 | | | |
| *Bifidobacterium* | | -0.47 | -1.54, 0.59 | 0.38 | | | |
| *Blautia/Anaerostipes* | | 0.68 | -0.19, 1.56 | 0.12 | *1.21* | *0.39, 2.02* | *0.004* |
| *Ruminococcus* | | -0.21 | -1.48, 1.06 | 0.74 | | | |
| **≥CR2** | | | | | | | |
| **Males** (ref: females) | 30 | 0.10 | -0.80, 1.00 | 0.82 | | | |
| **Age at vaccine** (years) | 30 | 0.04 | -0.10, 0.18 | 0.59 | | | |
| **Glucan initiated day 1** (ref: day 35) | 30 | -0.03 | -0.91, 0.85 | 0.94 | | | |
| **HCT** (ref: no HCT) | 30 | 0.06 | -1.00, 1.12 | 0.90 | | | |
| **MYCN amplified** (ref: non-amplified) | 30 | -0.30 | -1.25, 0.65 | 0.52 | | | |
| **α-diversity** | 30 | -0.04 | -0.10, 0.02 | 0.16 | | | |
| ***Number of prior relapses** (ref: 1)* | 30 | 0.19 | -0.57, 0.94 | 0.61 | | | |
| **Signatures** | 30 | | | | | | |
| *Ruminococcus* | | 1.03 | -0.86, 2.91 | 0.27 | | | |
| *Blautia/Faecalibacterium* | | -1.03 | -2.72, 0.67 | 0.23 | | | |
| *Romboutsia* | | -1.29 | -2.89, 0.32 | 0.11 | | | |
| *Blautia/Erysipelatoclostridium* | | 0.81 | -0.49, 2.11 | 0.21 | | | |
| *Bifidobacterium* | | 0.76 | -0.92, 2.44 | 0.36 | | | |
| *Blautia/Anerostipes* | | -0.91 | -3.04, 1.21 | 0.39 | | | |

*β coefficients and 95% confidence intervals were estimated using linear regression models of log-transformed anti-GD2 IgG1 titers, with separate models fitted for patients in CR1 and ≥CR2. Reference categories for categorical variables are indicated in parentheses. Variables with p<0.20 in univariable analyses were included in the corresponding multivariable models. Microbial signatures are derived from topic modeling. CI, confidence interval; CR, complete remission; HCT, hematopoietic cell transplantation.*

**Supplementary Table 7.** Cox regression analysis of PFS.

| | Univariable analysis | | | | Multivariable analysis | | |
|---|---|---|---|---|---|---|---|
| **Characteristic** | n | HR | 95% CI | p-val | HR | 95% CI | p-val |
| **CR1** | | | | | | | |
| **Males** (ref: females) | 75 | 0.91 | 0.33, 2.50 | 0.85 | | | |
| **Age at vaccine** (years) | 75 | 1.12 | 1.01, 1.24 | 0.01 | 0.97 | 0.82, 1.14 | 0.68 |
| **Glucan initiated day 1** (ref: day 35) | 75 | 1.56 | 0.55, 4.34 | 0.40 | | | |
| ***HCT*** *(ref: no HCT)* | 75 | 0.69 | 0.23, 2.02 | 0.50 | | | |
| **MYCN amplified** (ref: non-amplified) | 69 | 0.14 | 0.02, 1.10 | 0.06 | 0.14 | 0.02, 1.21 | 0.07 |
| ***α-diversity*** | *75* | 0.95 | 0.86, 1.04 | *0.28* | | | |
| **Signatures** | 75 | | | | | | |
| *Blautia/Romboutsia* | | 0.62 | 0.05, 7.56 | 0.71 | | | |
| *Erysipelatoclostridium* | | *16.74* | *3.37, 83.09* | *<0.001* | *20.73* | *2.05, 209.58* | *0.01* |
| *Faecalibacterium* | | 0.03 | 0.00, 8.74 | 0.23 | | | |
| *Bifidobacterium* | | 2.65 | 0.37, 18.77 | 0.33 | | | |
| *Blautia/Anaerostipes* | | 0.21 | 0.02, 2.74 | 0.23 | | | |
| *Ruminococcus* | | 0.11 | 0.00, 2.83 | 0.18 | 0.35 | 0.008, 15.51 | 0.59 |
| **≥CR2** | | | | | | | |
| **Males** (ref: females) | 38 | 0.93 | 0.38, 2.28 | 0.87 | | | |
| **Age at vaccine** (years) | 38 | 1.12 | 0.97, 1.30 | 0.14 | 1.10 | 0.95, 1.28 | 0.19 |
| **Glucan initiated day 1** (ref: day 35) | 38 | 0.76 | 0.29, 1.98 | 0.58 | | | |
| **HCT** (ref: no HCT) | 38 | 0.49 | 0.19, 1.24 | 0.13 | 0.45 | 0.17, 1.17 | 0.10 |
| **MYCN amplified** (ref: non-amplified) | 37 | 1.35 | 0.48, 3.77 | 0.57 | | | |
| ***Number of prior relapses*** | 38 | 1.05 | 0.52, 2.12 | 0.89 | | | |
| **α-diversity** | 38 | 1.02 | 0.96, 1.10 | 0.52 | | | |
| **Signatures** | 38 | | | | | | |
| *Ruminococcus* | | 0.41 | 0.05, 3.74 | 0.43 | | | |
| *Blautia/Faecalibacterium* | | 0.82 | 0.12, 5.80 | 0.85 | | | |
| *Romboutsia* | | 1.09 | 0.19, 6.11 | 0.92 | | | |
| *Blautia/Erysipelatoclostridium* | | 0.76 | 0.17, 3.37 | 0.72 | | | |
| *Bifidobacterium* | | 2.09 | 0.29, 15.16 | 0.46 | | | |
| *Blautia/Anerostipes* | | 2.13 | 0.32, 14.06 | 0.43 | | | |

*HRs and 95% confidence intervals were estimated using Cox proportional hazards regression models for progression-free survival, with separate models fitted for patients in CR1 and ≥CR2. Reference categories for categorical variables are indicated in parentheses. Variables with p<0.20 in univariable analyses were included in the corresponding multivariable models. Microbial signatures are derived from topic modeling. HR, hazard ratio; CI, confidence interval; CR, complete remission; HCT, hematopoietic cell transplantation.*

**Supplementary Table 8.** Cox regression analysis of progression-free survival in CR1 patients.

| | ***Univariable analysis*** | | | | ***Multivariable analysis*** | | |
|---|---|---|---|---|---|---|---|
| ***Characteristic*** | n | HR | 95% CI | p-val | HR | 95% CI | p-val |
| **Males** (ref: females) | 75 | 0.91 | 0.33, 2.50 | 0.85 | | | |
| **Age at vaccine** (years) | 75 | 1.12 | 1.01, 1.24 | 0.04 | 1.07 | 0.96, 1.21 | 0.19 |
| **Glucan initiated day 1** (ref: day 35) | 75 | 1.56 | 0.55, 4.38 | 0.40 | | | |
| ***HCT*** *(ref: no HCT)* | 75 | 0.69 | 0.23, 2.02 | 0.50 | | | |
| **MYCN amplified** (ref: non-amplified) | 69 | 0.14 | 0.02, 1.10 | 0.06 | 0.11 | 0.01, 0.95 | 0.045 |
| ***α-diversity*** | 75 | 0.95 | 0.86, 1.04 | 0.28 | | | |
| ***Clostridia*** *abundance* | *75* | *0.11* | *0.01, 0.96* | *0.046* | *0.03* | *0.003, 0.42* | *0.008* |

*HRs and 95% confidence intervals were estimated using Cox proportional hazards regression models for progression-free survival. Reference categories for categorical variables are indicated in parentheses. Variables with p<0.20 in univariable analyses were included in the corresponding multivariable models. HR, hazard ratio; CI, confidence interval; HCT, hematopoietic cell transplantation.*

**Supplementary Table 9**. Publicly available cohorts used as healthy microbiome references.

| *SRA/BioProject accession* | *Publication* |
|---:|---|
| *PRJNA695570* | Peterson et al. *Front. Microbiol*. 2021 |
| *PRJNA747761* | Kim et al. *ISME Commun*. 2022 |
| *PRJEB32762* | Soler-Saez et al. *mSystems* 2026 |
| *PRJNA706116* | De Filippis et al. *Nat Commun*. 2023 |
| *PRJEB24771* | Ordiz et al. *Am J Clin Nutr*. 2020 |
| *PRJEB48814* | *No associated publication* |

**Supplementary Table 10.** Cox regression analysis of progression-free survival.

| | Univariable analysis | | | |
|---|---|---|---|---|
| **Characteristic** | n | HR | 95% CI | p-val |
| **CR1** | | | | |
| **Males** (ref: females) | 64 | 1.14 | 0.40, 3.29 | 0.81 |
| **Glucan initiated day 1** (ref: day 35) | 64 | 1.47 | 0.51, 4.25 | 0.47 |
| **HCT** (ref: no HCT) | 64 | 0.73 | 0.24, 2.18 | 0.57 |
| **MYCN amplified** (ref: non-amplified) | 59 | NE* | - | - |
| **α-diversity** | 64 | 1.02 | 0.93, 1.12 | 0.68 |
| **High age gap** (ref: low) | 64 | *3.43* | *1.15, 10.28* | *0.028* |
| **≥CR2** | | | | |
| **Males** (ref: females) | 30 | 2.16 | 0.60, 7.76 | 0.24 |
| **Glucan initiated day 1** (ref: day 35) | 30 | 0.98 | 0.33, 2.94 | 0.97 |
| **HCT** (ref: no HCT) | 30 | 0.39 | 0.12, 1.28 | 0.12 |
| **MYCN amplified** (ref: non-amplified) | 30 | 1.33 | 0.37, 4.80 | 0.66 |
| ***Number of prior relapses*** | 30 | 1.25 | 0.35, 4.50 | 0.73 |
| **α-diversity** | 30 | 0.98 | 0.89, 1.08 | 0.71 |
| **High age gap** (ref: low) | 30 | 1.25 | 0.35, 4.50 | 0.73 |

*HRs and 95% confidence intervals were estimated using Cox proportional hazards regression models for progression-free survival, with separate models fitted for patients in CR1 and ≥CR2. Reference categories for categorical variables are indicated in parentheses. *For MYCN status in the CR1 cohort, the HR and 95% CI were not estimable using conventional Cox regression because all observed PFS events occurred in patients with non-amplified MYCN; therefore, the corresponding p-value could not be calculated. Multivariable models were not fitted because only one variable within each CR group met the prespecified criterion for inclusion (univariable p<0.20). HR, hazard ratio; CI, confidence interval; CR, complete remission; HCT, hematopoietic cell transplantation.*

**Supplementary Table 11.** Antibodies used for flow cytometry.

| ***Antigen*** | **Fluorochrome** | **Clone** | **Company** | **Catalog#** |
|---|---|---|---|---|
| *CD19* | APC | 1D3 | BD | 550992 |
| *CD3e* | BUV805 | 145-2C11 | BD | 749276 |
| *B220* | BUV615 | RA3-6B2 | BD | 569724 |
| *I-A/I-E (MHC-II)* | BUV737 | M5/114 | BD | 748845 |
| *CD45.2* | FITC | 104 | BD | 553772 |
| *CD8a* | BUV395 | 53-6.7 | BD | 563786 |
| *CD11c* | Brilliant Violet 421 | N418 | BD | 565451 |
| *CD11b* | BB700 | M1/70 | BD | 566417 |
| *NK1.1* | Alexa Fluor 700 | PK136 | BD | 560515 |
| *CXCR5* | PE | 2G8 | BD | 561988 |
| *CD4* | Brilliant Violet 650 | RM4-5 | BioLegend | 100545 |
| *F4/80* | Brilliant Violet 510 | BM8 | Biolegend | 123135 |
| *CD172a* | APC-Cy7 | P84 | Biolegend | 144018 |
| *XCR1* | Brilliant Violet 711 | ZET | Biolegend | 148245 |
